# HaloTag display enables quantitative single-particle characterization and functionalization of engineered extracellular vesicles

**DOI:** 10.1101/2023.09.25.559433

**Authors:** Roxana E. Mitrut, Devin M. Stranford, Beth N. DiBiase, Jonathan M. Chan, Matthew D. Bailey, Minrui Luo, Clare S. Harper, Thomas J. Meade, Muzhou Wang, Joshua N. Leonard

**Author notes:** These authors contributed equally as co-second authors (Devin M. Stranford, Beth N. DiBiase).

## Abstract

Extracellular vesicles (EVs) play key roles in diverse biological processes, transport biomolecules between cells, and have been engineered for therapeutic applications. A useful EV bioengineering strategy is to express engineered proteins on the EV surface to confer targeting, bioactivity, and other properties. Measuring how incorporation varies across a population of EVs is important for characterizing such materials and understanding their function, yet it remains challenging to quantitatively characterize the absolute number of engineered proteins incorporated at single-EV resolution. To address these needs, we developed a HaloTag-based characterization platform in which dyes or other synthetic species can be covalently and stoichiometrically attached to engineered proteins on the EV surface. To evaluate this system, we employed several orthogonal quantification methods, including flow cytometry and fluorescence microscopy, and found that HaloTag-mediated quantification is generally robust across EV analysis methods. We compared HaloTag-labeling to antibody-labeling of EVs using single vesicle flow cytometry, enabling us to measure the substantial degree to which antibody labeling can underestimate proteins present on an EV. Finally, we demonstrate the use of HaloTag to compare between protein designs for EV bioengineering. Overall, the HaloTag system is a useful EV characterization tool which complements and expands existing methods.

## INTRODUCTION

Extracellular vesicles (EVs) are nanoscale, lipid-encapsulated particles which natively transport biomolecular cargo between cells and play roles in diverse processes ranging from disease progression to wound healing.^1–5^ The most commonly studied EV populations, microvesicles (MVs) and exosomes, are produced via different cellular pathways—MVs form by direct budding from the cell’s plasma membrane, while exosomes originate by invagination within endosomal pathways to form multivesicular bodies that release exosomes by fusion to the cell surface.^3–5^ EVs are promising materials for therapeutic applications due to their ability to deliver functional cargo to recipient cells and their low immunogenicity and toxicity.^4–8^ While some EVs may naturally possess properties of therapeutic utility, here we focus on the distinct opportunities driven by a rapidly expanding suite of approaches for modifying EVs to confer new and useful properties.

One attractive strategy for conferring new functions to EVs is the addition of novel molecules to the EV surface through bioengineering. Display of native or synthetic biomolecular cargo molecules on EVs is commonly accomplished by genetically engineering cell lines to express a gene of interest, which is then actively or passively loaded into EVs during biogenesis.^8–12^ Targeting EVs to specific cell types has been widely investigated to improve therapeutic potential, typically by genetically fusing a targeting peptide, antibody, or antibody fragment to membrane proteins enriched in EVs. The earliest such report engineered EVs to display a rabies virus glycoprotein (RVG) peptide genetically fused to Lamp2b, a lysosome-associated membrane glycoprotein, conferring neuronal targeting mediated by RVG and gene knockdown when EVs are loaded with exogenous siRNA.^13^ We recently demonstrated that genetic display of antibody fragments on the EV surface confers enhanced cargo delivery to human T cells^10^. Genetically engineering EV producer cells has also been used to display biologics on EVs which exert therapeutic effects on recipient cells. For example, EVs loaded with truncated CD40 ligand, a transmembrane protein transiently expressed on activated T cells, fused to prostaglandin F2 receptor negative regulator (PTGFRN) induced more potent primary B cell activation compared to recombinant CD40 ligand.^14^ Surface-engineered EVs can also be used to investigate *in vivo* EV biodistribution and pharmacokinetics. For example, display of *Gaussia* luciferase by fusion to the C1C2 domains of lactadherin, which bind to phosphatidylserine on the EV surface, enables *in vivo* visualization and tracking after intravenous injection into mice,^15^ and a similar strategy can be used to track shedding of proteins from EVs^16^. Engineering EVs to bind albumin also directly increases their circulation time *in vivo*.^17^ As these examples illustrate, there now exists a substantial range of EV features that can be modulated by bioengineering the EV surface.

The EV surface can also be decorated post-biogenesis via physical modification. Representative examples of such methods include lipid tag-driven insertion of cargo into the EV membrane, EV fusion with liposomes, or chemical modification wherein cargo is covalently linked to functional groups on EV membrane proteins.^8,9,12,18–20^ For example, EV surface-display of cell penetrating peptides can increase EV uptake by inducing micropinocytosis; such functionalization has been achieved by techniques including conjugation of the peptide to lipids followed by lipid insertion or by covalent conjugation of the peptide directly to the EV surface.^21–23^ Lipid insertion has also been used to decorate EVs with epidermal growth factor receptor (EGFR)-binding nanobodies and PEG, enabling EV-targeting to EGFR-expressing cells *in vitro* and increased circulation time *in vivo* due to PEG shielding.^19^ In addition to biologics, post-biogenesis modification can enable the addition of synthetic molecules to EVs, such as quantum dots (QD) which enable high-resolution live-imaging microscopy.^24^ Although post-biogenesis EV modification requires careful consideration of scale up and downstream purification, these methods are increasingly feasible for implementing a range of EV modifications.

The growing availability and utility of methods for EV surface functionalization have highlighted the perhaps counterintuitive observation that it remains difficult to quantify surface proteins on EVs in absolute terms.^25,26^ Many common EV protein quantification techniques, such as ELISA, semi-quantitative Western blots, single vesicle flow cytometry, and single particle imaging systems such as ExoView (NanoView Biosciences), rely on antibody binding to EV surface markers. These methods enable relative quantification of the protein/s of interest. However, absolute quantification is generally not possible with these techniques. One challenge is that each antibody’s labeling efficiency is inherently linked to the specificity and affinity of the antibody to the protein of interest, which can vary between antibody target, antibody type, and even antibody lot. A particular challenge for quantifying high copy number proteins is that antibody-mediated detection may be limited by crowding (steric occlusion) as antibodies accumulate on the EV surface, causing antibody labeling to saturate before all target sites are bound. While bulk EV measurements may overcome some such limitations associated with the native EV structure, only single vesicle measurements can ultimately provide insight into key properties including the distribution (and associated heterogeneity) of protein display across a population of EVs. Fluorescence microscopy has recently been used to measure single EVs and directly quantify fluorescent protein cargo loaded into EVs (in this case, avoiding the need for antibody labeling).^27^ This technique is limited by the number of vesicles that can be readily measured in a typical experiment, which is frequently in the tens of vesicles. Single vesicle-flow cytometry (SV-FC) enables analyzing individual particles as small as 100 nm and has been used to investigate EV properties at a single EV resolution, and it is useful for detecting fluorescent proteins and synthetic dyes.^14,27–30^ Across these methods, it remains challenging to measure low numbers of proteins per EV due to the relative low quantum yield of fluorescent proteins (i.e., compared to synthetic dyes). Altogether these observations identify an opportunity for complementing existing approaches by developing methods for absolute quantification of EV surface proteins at the single EV level; pursuing this goal motivates this study.

Here we investigate a versatile strategy for quantifying engineered surface proteins on EVs using the HaloTag system. HaloTag is a modified bacterial haloalkane dehalogenase that has been mutated to rapidly, irreversibly, and under mild conditions form a covalent bond with a specific chloroalkane HaloTag ligand.^31–34^ We found that display of HaloTag protein on EVs enables surface functionalization via reaction with HaloTag ligands, enabling quantitative single-particle characterization of engineered proteins on EVs. We compare such analyses across a range of quantification methods. The reagents and methods described here will contribute to the growing suite of tools enabling rigorous EV analysis.

## MATERIALS AND METHODS

### General DNA assembly

Plasmids used in this study were generated using standard polymerase chain reaction techniques and type II and IIs restriction enzyme cloning. Restriction enzymes, Phusion DNA polymerase, T4 DNA Ligase, and Antarctic phosphatase were purchased from New England Biolabs. psPAX2 and pMD2.G plasmids were gifted by William Miller from Northwestern University. HaloTag, a gift from Veit Hornung (Addgene, plasmid #80960),^35^ was genetically fused at the C-terminus to a validated PDGFR transmembrane domain^36^ and was cloned into a pGIPZ lentiviral expression vector (Open Biosystems) using PacI and AgeI restriction enzymes. The novel PTGFRN transmembrane domain display systems were codon optimized and DNA synthesized by Thermo Fisher, then cloned into the HaloTag plasmid using SapI and NotI restriction enzymes. The HA-tag HaloTag plasmid was cloned using PCR to replace the N-terminal FLAG-tag with the HA-tag sequence. All plasmids were sequence-verified. Plasmids were transformed into TOP10 competent *E. coli* and grown at 37°C.

### Plasmid preparation

Plasmid DNA used to generate lentivirus for cell line engineering was prepared using a polyethylene glycol precipitation protocol.^37^ Plasmid DNA used for transient transfection of HEK293FT cells was prepared using a MidiPrep protocol (Omega Bio-Tek, D6904-04). DNA purity and concentrations for relevant experiments were measured with a NanoDrop 2000 (Thermo Fisher Scientific).

### Cell culture

HEK293FT cells (Thermo Fisher, R70007) and sublines generated from this line were cultured in Dulbecco’s Modified Eagle Medium (DMEM, Gibco 31600-091) supplemented with 10% FBS (Gibco, 16140-071), 1% penicillin-streptomycin (Gibco, 15140-122), and 4 mM additional L-glutamine (Gibco, 25030-081). Cells were subcultured at a 1:5 or 1:10 ratio every 2-3 d, using Trypsin-EDTA (Gibco, 25300-054) to remove adherent cells from the plate. Cells were maintained at 37°C and 5% CO_2_.

### Vector and stable cell line generation

HEK293FT cells were used to produce lentivirus for stable cell line generation. HEK293FT cells were plated in 10 cm dishes at a density of 5 x 10^6^ cells/dish. Cells were transfected 7 h later with 10 μg of viral transfer vector, 8 μg psPAX2, and 3 μg pMD2.G via calcium phosphate transfection.^37^ Cell culture medium was changed 12-16 h later. Lentivirus was harvested from the conditioned medium 28 h post media change and centrifuged at 500 g for 2 min to clear cells. The supernatant was filtered through a 0.45 μm pore filter (VWR, 76479-020). Lentivirus was concentrated from the filtered supernatant by ultracentrifugation in Ultra Clear tubes (Beckman Coulter, 344059) at 100,420 g at 4°C for 90 min in a Beckman Coulter Optima L-80 XP ultracentrifuge using an SW41Ti rotor. Lentivirus was stored on ice until use. 1 x 10^5^ cells were plated 24 h before transduction in a 12-well plate. At the time of transduction, cell culture medium was aspirated, and lentivirus was added. Drug selection began 2 d post transduction with 1 μg/mL puromycin (Invitrogen, ant-pr-1). Cells were kept in antibiotics for at least two weeks with subculturing every one to two days before further characterization.

### EV production and isolation

HEK293FT cells were plated in 15 cm dishes at a density of 15 x 10^6^ cells/dish. The following morning, the medium was replaced with DMEM supplemented with 10% EV-depleted FBS (Gibco, A2720801). After 24 h, the conditioned medium was harvested as previously reported and the EV populations were separated via sequential centrifugation.^38^ Cell debris and apoptotic bodies were removed by centrifugation spins at 300 g for 10 min and 2,000 g for 20 min, respectively. Microvesicles were pelleted at 15,000 g for 30 min in a Beckman Coulter Avanti J-26XP centrifuge using a J-LITE JLA 16.25 rotor. Exosomes were pelleted at 120,417 g for 135 min in a Beckman Coulter Optima L-80 XP model using a SW41 Ti rotor. All centrifugation was performed at 4°C. EVs were resuspended in residual medium remaining in their respective vessel following supernatant removal. EVs were characterized according to MISEV 2018 guidelines, as described in Results.^39^

### Nanoparticle tracking analysis (NTA)

Vesicle concentration and size were measured using a Nanosight NS300 (Malvern) running software v3.4 and a 642 nm laser. Vesicles were diluted to 2-10 x 10^8^ particles/mL in phosphate buffered saline (PBS) for analysis. Samples were run at an injection rate of 30, imaged at a camera level of 14, and analyzed at a detection threshold of 7. Three 30 second videos were captured for each sample to determine the average vesicle concentration and size histograms.

### HaloTag ligand conjugation and purification

HaloTag ligand conjugation was performed following manufacturer recommendation for cells. 10^9–10^ EVs per sample were adjusted to equivalent concentrations and volumes (set by the lowest yield sample in any one experiment) with EV-depleted DMEM, in black 1.7 mL microcentrifuge tubes to decrease dye photobleaching. HaloTag ligand was diluted 1:200 from the manufacturer stock with EV-depleted DMEM and added to each tube so that it made up 1/5 of the total volume. The reaction was run at 37°C for 15 min. Free dye was removed via 30 kDa centrifugal spin filters (MilliporeSigma, UFC9030). Three wash spins were conducted per sample, where 5 mL of PBS was added and then samples were spun at 5,000 g for 5 min, resulting in 300-500 μL of conjugated EVs. A dye-only sample that we term “Mock” included no EVs and was reacted under the same conditions as the EV samples; this control was included to evaluate any free dye remaining after washing. NTA was again performed to determine EV concentrations.

### Gold nanoparticle synthesis and conjugation

Gold nanoparticle (AuNP) synthesis is described in detail in the **Supplementary Information**. Briefly, citrate-stabilized AuNPs with 12.3 ± 1.3 nm core diameter were synthesized and characterized by TEM and UV/vis (**Figure S1**). Coupling-ready HaloTag ligand was synthesized and characterized by NMR (**Scheme S1, Figure S2**). Short, cysteine-terminated peptide chains were synthesized (**Scheme S2, Figure S3**) and used to replace citrate on the AuNP surface with HaloTag ligands (**Figure S4**), resulting in functionalized AuNPs for HaloTag immunogold labeling.

For AuNP HaloTag ligand conjugation, 10^9^ EVs per sample were adjusted to equivalent concentrations and volumes with PBS. HaloTag AuNP ligand stock (0.7 μM) was added to each tube so that it made up 1/5 of the total volume. The reaction was run at room temperature (approximately 20°C) for 16 h, and the resulting reactions were used without further purification.

### Transmission electron microscopy (TEM)

10 μL of purified vesicles was placed onto a carbon-coated copper grid (Electron Microscopy Services, CF400-Cu-50) for 10 min before excess liquid was wicked away with a piece of filter paper. The grid was dipped in PBS twice to remove excess proteins and unreacted ligands from the media and reaction, and was allowed to dry for 2 min. To achieve negative staining, 10 μL of uranyl acetate solution (2 wt% in Milli-Q water) was placed on the grid for 1 min before being wicked away with filter paper. The grid was allowed to fully dry (3 h to overnight) at room temperature (approximately 20°C). Bright-field TEM imaging was performed on a JEOL 1230 TEM. The TEM operated at an acceleration voltage of 100 kV. All TEM images were recorded by a Hamamatsu ORCA side-mounted camera or a Gatan 831 bottom-mounted CCD camera, using AMT imaging software. TEM images were analyzed using ImageJ (Fiji 370).^40^

### Fluorescence microscopy

Fluorescence microscopy measurements were performed using a custom modified Olympus IX73 inverted microscope equipped with optical components from ThorLabs unless otherwise stated. All optical filters were acquired from Semrock. Alexa Fluor 488 HaloTag ligand (Promega, G1001) was excited by a 473-nm laser (Laser Quantum, 500 mW), which was passed through a bandpass filter (LD01-473/10– 12.5), a half-wave plate, and a quarter-wave plate. The intensity of the 473-nm laser was 0.75 kW/cm2. Videos of the EVs and Alexa Fluor-functionalized beads were acquired with an exposure time of 5 ms over a 32 x 32 µm area. Samples were prepared on Piranha-treated #1.5 glass coverslips. Piranha solution consisted of H_2_SO_4_ and 30% H_2_O_2_ solution at a 3:1 ratio by volume, and the coverslips were treated for at least 3 h. Poly(l-lysine) (PLL) solution (0.1 mg/mL in water) was deposited onto coverslips to provide an adhesive surface for vesicles. Suspensions containing vesicles were drop casted directly and were allowed to completely dry. Microscope images showed vesicles to be well spaced, limiting the interference between close features. Features were chosen manually based on size and brightness, and ImageJ was used to determine fluorescence intensity, which was corrected for background fluorescence. A minimum of 300 events were analyzed for fluorescence intensity.

### Fluorescent protein SDS-PAGE gel quantification of HaloTag-loading on EVs

To label EVs, 10^9^ vesicles were incubated with 2 µL of 1:200 diluted HaloTag ligand at 37°C for 15 min. Fluorescent Compatible Sample Buffer (Thermo Fisher, LC2570) with 100 mM β-mercaptoethanol (BME) was added to the samples followed by incubation at 70°C for 3 min. Each sample was run in duplicate, with 4.5 x 10^8^ EVs/lane, on a 4–15% Mini-PROTEAN® TGX™ Precast Protein Gels (BioRad, 4561086) and run at 100 V for 1 h. To quantify HaloTag protein loading on EVs, several concentrations of purified recombinant HaloTag protein standard (Promega, G4491) were labeled with ligand and run in duplicate to generate a calibration curve. Samples and the calibration curve were run on the same protein gel or on multiple protein gels which were imaged together to enabling direct comparison of fluorescence intensity. Gels were washed in Milli-Q water and imaged using an Azure Sapphire Imager. Gel fluorescent images were analyzed using ImageJ (Fiji 370).

### Cell lysate generation

HEK293FT cell lines were washed with cold (∼4°C) PBS and lysed with ice-cold radioimmunoprecipitation assay buffer (150mm NaCl, 50mm Tris-HCl pH 8.0, 1% Triton X-100, 0.5% sodium deoxycholate, 0.1% sodium dodecyl sulfate) supplemented with protease inhibitor (Pierce/Thermo Fisher, #A32953). After a 30 min incubation on ice, lysates were centrifuged at 14,000 g for 20 min at 4°C. Protein concentration for each sample was evaluated using a bicinchoninic acid (BCA) assay (Pierce/Thermo Fisher, #23225). Samples were kept on ice until use or frozen at −80°C for long term storage.

### Western blotting cell lysate and EVs

For western blots comparing protein in cell lysates and protein in vesicles, a fixed number of vesicles and a fixed mass of cell lysate were loaded into each well: 4.5 x 10^8^ EVs/lane and 2 μg protein/lane, respectively. An established western blot protocol^37^ was followed with the following modifications. In most cases, the following reducing Laemmli composition was used to boil samples (60 mM Tris-HCl pH 6.8, 10% glycerol, 2% sodium dodecylsulfate, 100 mM dithiothreitol (DTT), and 0.01% bromophenol blue); in some cases, a nonreducing Laemmli composition (without DTT) was used, as previously reported.^41^ After transfer, membranes were blocked while rocking for 1 h at room temperature in 5% milk in Tris-buffered saline with Tween (TBST) (pH: 7.6, 50mm Tris, 150mm NaCl, HCl to pH 7.6, 0.1% Tween 20). Primary antibody was added in 5% milk in TBST, rocking, for 1 h at room temperature and then washed three times with TBST for 5 min each. Secondary antibody in 5% milk in TBST was added at room temperature for 1 h or overnight at 4°C. Membranes were then washed three times with TBST for 5 min each. The membrane was incubated with Clarity Western ECL substrate (Bio-Rad, #1705061) and imaged on an Azure c280 running Azure cSeries Acquisition software v1.9.5.0606. Specific antibodies, antibody dilution, heating temperature, heating time, and Laemmli composition for each antibody was used as previously reported.^41^

### Surface staining cells and EVs

For surface staining cells, medium was aspirated, and cells were harvested with 1 mL cold (4°C). fluorescence-activated cell sorting (FACS) buffer (PBS pH 7.4, 2mM EDTA, 0.05% bovine serum albumin). Samples were centrifuged at 150 g for 5 min at 4°C. The supernatant was removed, and cells were resuspended in FACS buffer (50 μL) and blocked with 10 μL of 1 mg/mL IgG (Thermo Fisher, Human IgG Isotype Control, 02-7102, RRID: AB_2532958) for 5 min at 4°C. Next, FLAG-tag antibody (Bio-Techne, DYKDDDDK Epitope Tag Alexa Fluor® 488-conjugated Antibody, IC8529G) or HA-tag antibody (BioLegend, Alexa Fluor® 488 anti-HA.11 Epitope Tag Antibody, 901509) was added at a concentration of 1 µg/10^6^ cells or 0.5 µg/10^6^ cells, respectively, and cells were incubated at 4°C for 30 min. Cells were washed three times by adding cold FACS buffer (1 mL), centrifuging cells at 150 g for 5 minutes at 4°C, and decanting supernatant. Cells were resuspended in 1-2 drops of FACS buffer prior to analytical flow cytometry. For surface staining EVs, 3 x 10^8^ EVs were adjusted to a total volume of 20 µL with PBS. EVs were blocked with 1 µL of 1 mg/mL IgG for 10 min on ice, after which 1 µg of the FLAG-tag antibody or 0.1 µg of the HA-tag antibody was added and incubated for 45 min on ice. PBS was added to bring the final volume to 250 µL. The FLAG-tag antibody was spun at 14,000 g for 1 h at 4°C prior to use (to pellet and remove any aggregates).

### EV adsorption onto latex beads

For analyzing bulk EV properties, 10^9^ EVs per sample were adjusted to the equivalent concentrations and volumes with PBS. 2 µL of aldehyde/sulfate latex beads (Thermo Fisher, A37304) diluted in PBS 1:10 were added to the samples and incubated for 15 min at room temperature. PBS was added to bring the final volume to 200 µL, and samples were rocked for 2 h at room temperature. Samples were used immediately or stored at 4°C overnight.

### Analytical flow cytometry and analysis of cells and beads

Flow cytometry was performed on a BD LSR Fortessa Special Order Research Product. To detect AF 488, a 488 nm laser with a 505 nm long pass filter and a 530/30 nm bandpass filter were used. To detect AF 660, a 552 nm laser with a 600 nm long pass filter and a 610/20 nm bandpass filter were used. Approximately 10,000 live cells or latex beads were collected per sample for analysis. Data were analyzed using FlowJo v10 (FlowJo, LLC) as described in detail in the **Supplementary Information** (**Figures S5-6**). Briefly, cells were identified using an FSC-A vs SSC-A plot and gated for singlets using an FSC-A vs FSC-H plot. Mean fluorescence intensity (MFI) of single-cell samples was exported and averaged across three biological replicates. Autofluorescence from untreated cells was subtracted from other samples. Latex beads were identified using an FSC-A vs SSC-A plot. Mean fluorescence intensity (MFI) of bead samples was exported and averaged across three technical replicates. Autofluorescence from untreated EV samples adsorbed onto beads was subtracted from other samples. Standard error of the mean was propagated through calculations.

### Analytical flow cytometry and analysis of EVs

Single vesicle flow cytometry was performed on either an Apogee Micro Plus vesicle flow cytometer, NanoFCM Flow Nanoanalyzer vesicle flow cytometer, or on a BD LSR Fortessa Special Order Research Product. On the Apogee Micro Plus, a 488 nm laser was used with a 530/40 nm bandpass filter. For each sample, 3 x 10^8^ EVs were adjusted to a total volume of 250 µL using PBS and loaded onto a black 96-well plate to minimize dye photobleaching. Samples were run at 1.5 µL/min for 1 min. Quantification beads were run at 10.5 µL/min for 1 min. Data were analyzed using FlowJo v10 (FlowJo, LLC) as described in detail in the **Supplementary Information** (**Figure S7**). Briefly, EVs were identified using a 405-LALS(Area) vs 405-SALS(Area) plot and gated as EVs using a 488-Grn(Peak) vs 405-LALS(Peak) plot. Mean fluorescence intensity (MFI) of EV samples was exported and averaged across three technical replicates. Autofluorescence from unmodified HEK293FT EVs was subtracted from other samples to identify the fluorescent signal attributable to the dye. Standard error of the mean was propagated through calculations.

On the NanoFCM Flow Nanoanalyzer, a 488 nm laser was used with a 488/10 nm (for SSC) and 525/40 nm (for AF 488) bandpass filters, and a 640 nm laser was used with a 710/40 nm bandpass filter (for AF 660). For each sample, 2 x10^8^ or 1 x10^9^ EVs were adjusted to a total volume of 100 µL using 0.1 µm filtered PBS and loaded into microtubes (Axygen, MCT-060-C). After instrument start up, Quality Control (QC) Beads (250 nm Fluorescent Silica Microspheres) were run at a 100x dilution with large particle thresholding for alignment, focusing, and calibration. Following QC checks, a Silica Nanosphere Cocktail (S16M-Exo) was run at a 100x dilution with small particle thresholding for use as a size standard to create a calibration curve of particle size and side scatter intensity. After setup and calibration, samples were run at the NanoFCM software standard sample flow rate, as determined by the sampling pressure of 1.0 kPa, and recorded for 2 min. After data collection, samples were pre-analyzed using the NanoFCM software. Briefly, each sample was thresholded on small particles, and samples were background subtracted using the “Set Blank” software feature, where the blanks were PBS containing media and HaloTag ligand or antibody in the same dilution as their corresponding samples. Data were analyzed using FlowJo v10 (FlowJo, LLC) as described in detail in the **Supplementary Information** (**Figure S8**). Mean fluorescence intensity (MFI) of EV samples was exported and averaged across three technical replicates. Fluorescence from unmodified HEK293FT EVs labeled with HaloTag ligand or antibody was subtracted from other samples to identify the fluorescent signal attributable to the EV labeling. Standard error of the mean was propagated through calculations.

On the BD LSR Fortessa Special Order Research Product, a 488 nm laser was used with a 505 nm long pass filter and a 530/30 nm bandpass filter to detect AF 488 ligand, and a 552 nm laser was used with a 600 nm long pass filter and a 610/20 nm bandpass filter to detect AF 660. After conjugation with HaloTag ligand, samples were washed with 0.1 µm filtered PBS to minimize non-EV events detected, and samples were then run on low speed for 1 min. A threshold was applied on SSC-H to minimize very small non-EV events. Data were analyzed using FlowJo v10 (FlowJo, LLC) as described in detail in the **Supplementary Information** (**Figure S9**). Briefly, small, non-fluorescent non-EV background events were identified using a sample of 0.1 µm filtered PBS on a FITC-H vs SSC-A plot for AF 488 conjugated EVs and on a PE Texas Red-H vs SSC-A plot for AF 660 conjugated EVs. EVs in the size range of interest (<590 nm) were identified using ApogeeMix Size reference beads (Apogee, 1527) on a FITC-H vs SSC-A or PE Texas Red-H vs SSC-A plot. Mean fluorescence intensity (MFI) of EV samples was exported and averaged across three technical replicates. Autofluorescence from unmodified HEK293FT EVs (i.e., not loaded with HaloTag) that were mixed with dye and washed was subtracted from other EV samples to identify the fluorescent signal attributable to HaloTag-mediated conjugation of dye. Standard error of the mean was propagated through calculations.

### Quantification of HaloTag-conjugation on EVs from flow cytometry data

To determine the absolute quantity of HaloTag ligands conjugated to EVs, quantification beads for AlexaFluor 488 (Bangs Lab, Quantum™ Alexa Fluor® 488 MESF, 488) were run on the Apogee or BD flow cytometer for each experiment to enable calibration of fluorescent signal in absolute units. One drop of each fluorescent and blank bead was added to 250 µL of PBS. Data were analyzed using FlowJo v10 (FlowJo, LLC) as described in detail in the **Supplementary Information** (**Figure S10**). Briefly, beads were identified using a 488-Grn(Peak) vs 405-LALS(Peak) plot. Autofluorescence of the blank bead was subtracted from the other beads. A linear regression of the measured bead fluorescence versus the known number of Alexa Fluor 488 in each bead (as reported by the manufacturer) was calculated, and the resulting quantification curve was used to convert EV mean fluorescence intensity (MFI) into absolute Alexa Fluor 488 units (MESF).

### Statistical analysis

Statistical tests employed are described in relevant figure legends. Unless otherwise stated, three independent biological replicates (cells) or technical replicates (beads and EVs) were analyzed per condition, and the mean fluorescence intensity of approximately 10,000 live single cells or beads, or 200,000 EVs were analyzed per sample. Unless otherwise indicated, error bars represent the standard error of the mean. GraphPad Prism 9.2 was used to analyze the data. Pairwise comparisons were made using unpaired Student’s t-test. Multicomparison statistical analysis was performed using a one-way ANOVA test, followed by Tukey’s multiple comparison (i.e., honestly significant difference, HSD) test to evaluate specific comparisons. Significance threshold: *p < 0.05, **p < 0.01, ***p < 0.001, ****p < 0.0001.

## RESULTS

### Engineered EVs display functional HaloTag on their surface

In order to investigate whether the HaloTag system can be used to evaluate EV surface display, we first genetically fused the HaloTag gene to a platelet-derived growth factor receptor transmembrane domain (PDGFR-TMD) using a strategy that previously has led to display of various domains onto EV membranes.^36,38,42,43^ Lentiviral vectors were used to stably express the resulting fusion protein in human embryonic kidney (HEK293FT) EV producer cells (**Figure 1A**).^44^ EVs were then harvested from producer cells by differential centrifugation to remove cell debris and apoptotic bodies, resulting in enriched exosome (120,417 g) and microvesicle (MV, 15,000 g) fractions; throughout this study, these terms are defined by the separation applied to generate each EV fraction.^38^ EVs were then characterized according to MISEV 2018 guidelines to ensure that the samples isolated exhibit expected size, shape and protein markers.^39^ EVs exhibited typical cup-shape morphology under Transmission Electron Microscopy (TEM) (**Figure 1B**). Western blots for detecting common EV markers confirmed the presence of ALIX, CD9 and CD81 in both fractions, while calnexin—an endoplasmic reticulum marker—was only present in cell lysate (**Figure 1C-D**). Expression of the HaloTag fusion protein in cell lysate and in exosomes and MVs was confirmed by western blot (**Figure 1D**). Nanoparticle tracking analysis revealed that particles generated from HaloTag-expressing cells were ∼100-200 nm in size, consistent with previously reported HEK293FT EV sizes (**Figure 1E**).^45^

**Figure 1:**
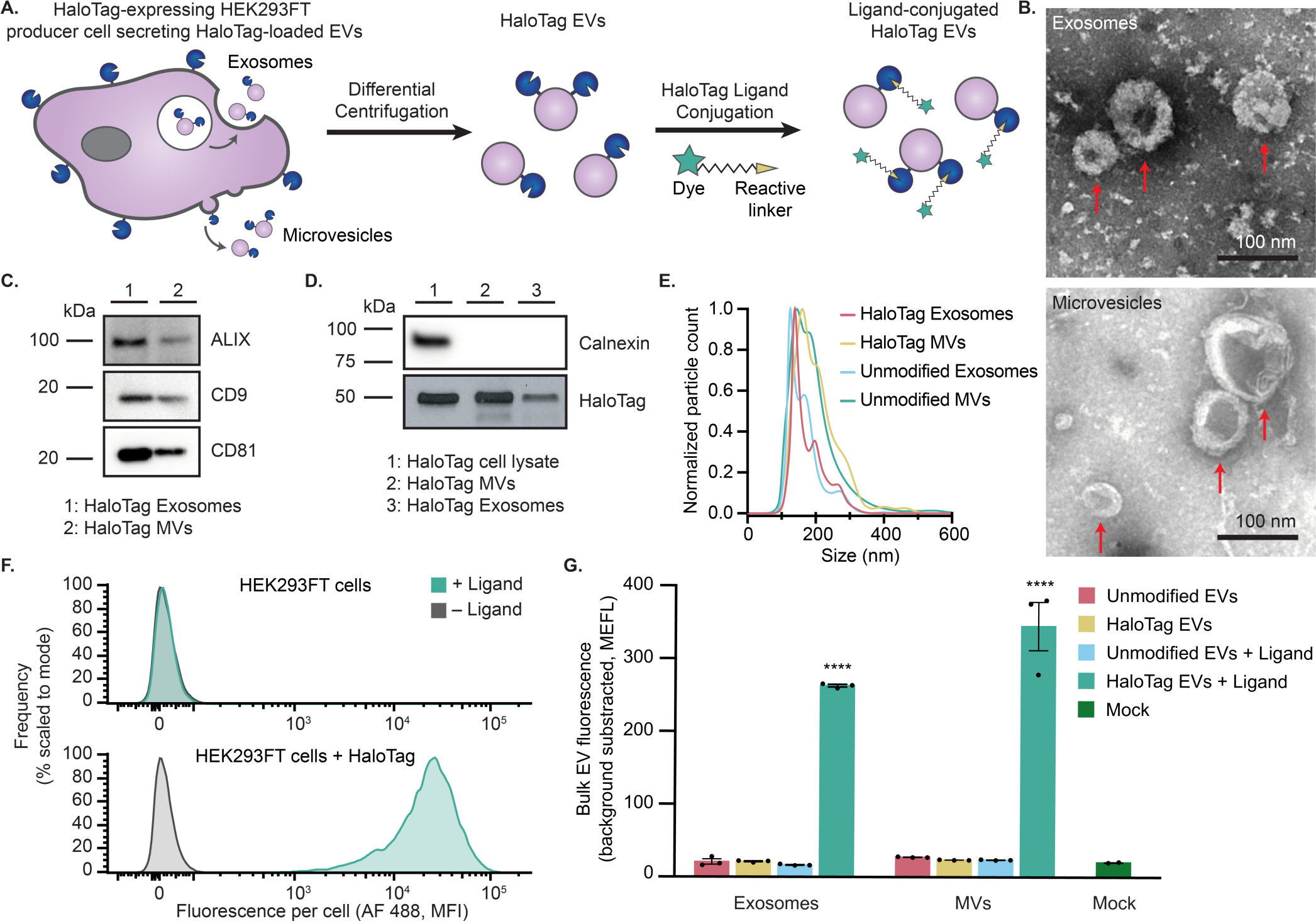
Validation of a HaloTag display system for functionalizing the EV surface. A) Cartoon illustrating how HaloTag-expressing HEK293FT cells produce HaloTag EVs which are isolated via differential centrifugation. The HaloTag protein can be used to display a variety of moieties on the surface of EVs via conjugation with different HaloTag ligands. B) HaloTag EVs display classical EV morphology. Transmission electron micrographs of HaloTag EV subpopulations show classic cup-shaped morphology. Top: Exosomes. Bottom: MVs. Scale bar: 100 nm. C,D) Western blots yield expected patterns of common EV markers in purified vesicles vs producer cells. EVs contain expected markers, calnexin is only present in cell lysate, and the FLAG tag fused to N-terminus of engineered HaloTag yields an expected band of 44.8 kDa and is present in engineered cell lysate and EVs populations. E) Representative histogram of nanoparticle tracking analysis of EVs derived from HEK293FT cells with or without HaloTag expression, normalized to the mode in each population. F) HaloTag expressing cells (bottom) but not unmodified cells (top) increase in fluorescence after exposure to AF 488 HaloTag ligand. G) HaloTag EVs adsorbed on polystyrene beads react with and conjugate AF 488 ligand, but unmodified EVs and the mock condition yield no such signal. Cell experiments were performed in biological triplicate. EV experiments were performed in technical triplicate. Error bars indicate standard error of the mean. Multicomparison statistical analysis was performed using a one-way ANOVA test, followed by Tukey’s multiple comparison test to evaluate specific comparisons (*p < 0.05, **p < 0.01, ***p < 0.001, ****p < 0.0001).

To evaluate whether the HaloTag was functional and displayed on the surface membrane in the correct orientation, we first reacted a HaloTag ligand with the parental EV-producer cells. A cell-impermeable HaloTag ligand coupled to Alexa Fluor 488 (AF 488) was selected so that increases in cell and EV fluorescence could be specifically attributed to ligand reaction with extracellular-facing HaloTag constructs. HaloTag-expressing cells showed a ∼1000-fold increase in fluorescence when reacted with ligand, whereas control cells did not increase in fluorescence upon exposure to ligand, demonstrating that there were no detectable non-specific ligand-cell interactions and that post-reaction washing steps were sufficient to remove excess unreacted or free ligand (**Figure 1F**). A similar trend was also observed in bulk EV properties; EVs (adsorbed onto latex beads prior to flow cytometry) demonstrated a clear increase in fluorescence in a manner that depended upon both HaloTag expression and ligand addition (**Figure 1G**).

### HaloTag labeling enables absolute quantification of EV display at single vesicle resolution

Single vesicle-flow cytometry (SV-FC) is an emerging method capable of analyzing individual particles as small as 100 nm, and it enables investigating EV properties at a single EV resolution.^28,29,46–48^ Notably, properties pertaining to the distribution of EV states are uniquely evaluable using such single particle methods. Using an Apogee Micro flow cytometer (**Figure 2A**), we observed that as many as 90% of HaloTag EVs were successfully labeled with HaloTag ligand, demonstrating that a majority of the engineered EVs contained the HaloTag construct (**Figure 2B**). Fluorescent quantification beads can be used to calculate the absolute number of AF 488 fluorophores conjugated to each HaloTag EV.^28,49^ The fluorescently-labeled EVs exhibit means as high as ∼1,500 HaloTag protein constructs per EV on both exosomes and MVs (**Figure 2C**). To evaluate whether the HaloTag ligand conjugation reaction proceeded to completion to yield fully-labeled EVs, we varied ligand concentration and reaction time and observed minimal changes in labeling, indicating that the HaloTag labeling was saturated (**Figure S11-S12**). These single vesicle measurements agree with bulk EV observations that HaloTag labeling of EVs is specific and has no detectable background fluorescence, while providing novel insight into heterogeneity in protein display across the EV population.

**Figure 2:**
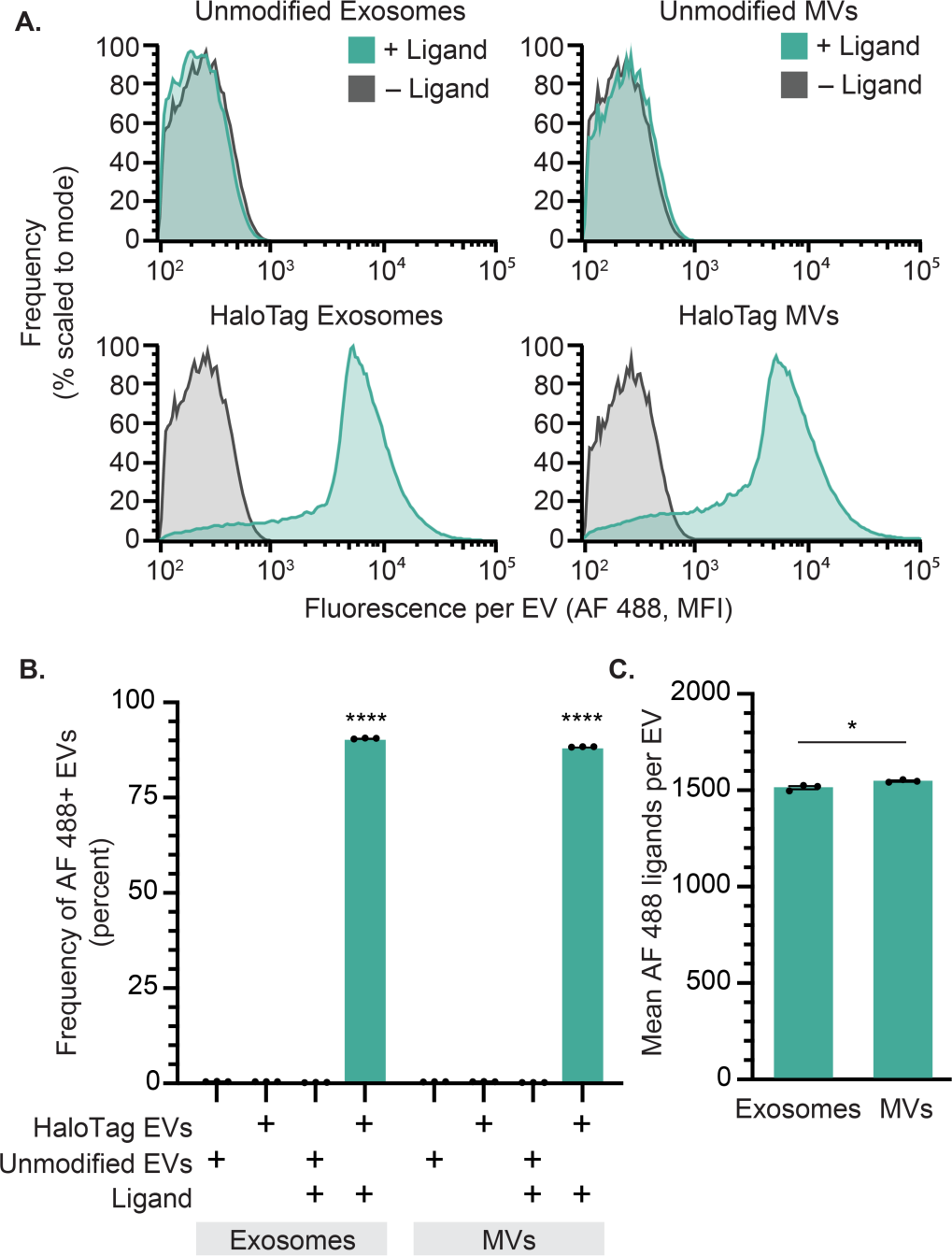
HaloTag enables quantification of surface display at a single vesicle resolution. A) HaloTag-displaying exosomes and MVs (bottom) but not unmodified exosomes and MVs (top) increase in fluorescence upon reaction with AF 488 ligand, measured by Apogee SV-FC. B) HaloTag labeling of exosomes and MVs is efficient and specific (i.e., background is very low). C) The average number of AF 488 ligands per EV can be determined using calibration based upon quantification beads (**Figure S10)**. All experiments were performed in technical triplicate, and error bars indicate standard error of the mean. Multicomparison statistical analysis was performed using a one-way ANOVA test, followed by Tukey’s multiple comparison test to evaluate specific comparisons (*p < 0.05, **p < 0.01, ***p < 0.001, ****p < 0.0001).

### HaloTag EV surface multi-functionalization can be titrated

A potential utility of HaloTag labeling, facilitated by the high level of HaloTag surface display achieved by the PDGFR-TMD system, is the ability to install multiple ligand types per particle to create multifunctional EVs (**Figure 3A**). To investigate this possibility using our dye-based model system, we varied the relative amounts of two different HaloTag ligands in the conjugation reaction. Alongside our initial AF 488 ligand, we used a second cell-impermeable Alexa Fluor 660 (AF 660) ligand, as these fluorophores can be spectrally separated. Per manufacturer descriptions, the ligands have different reaction rates, and thus ligand conjugation was performed at their recommended concentrations (AF 488 was reacted at 1 µM, AF 660 was reacted at 3.5 µM). Since the Apogee SV-FC system used in this study does not have a laser which can detect AF 660 directly, this initial assay was designed to quantify co-labeling as a decrease in AF 488 fluorescence. AF 488 fluorescent labeling of EVs decreased with increasing AF 660 ligand ratios (**Figure 3B**). The average number of AF 488 ligands measured per EV also decreased as the AF 660 ligand concentration increased (**Figure 3C**), showing a linear relationship for both EV populations when accounting for the relative ligand reaction rates (**Figure 3D**). Thus, using the AF 488 fluorescence as a proxy, we can infer co-labeling of HaloTag and can tune relative conjugation extents by adjusting relative ligand concentrations.

**Figure 3:**
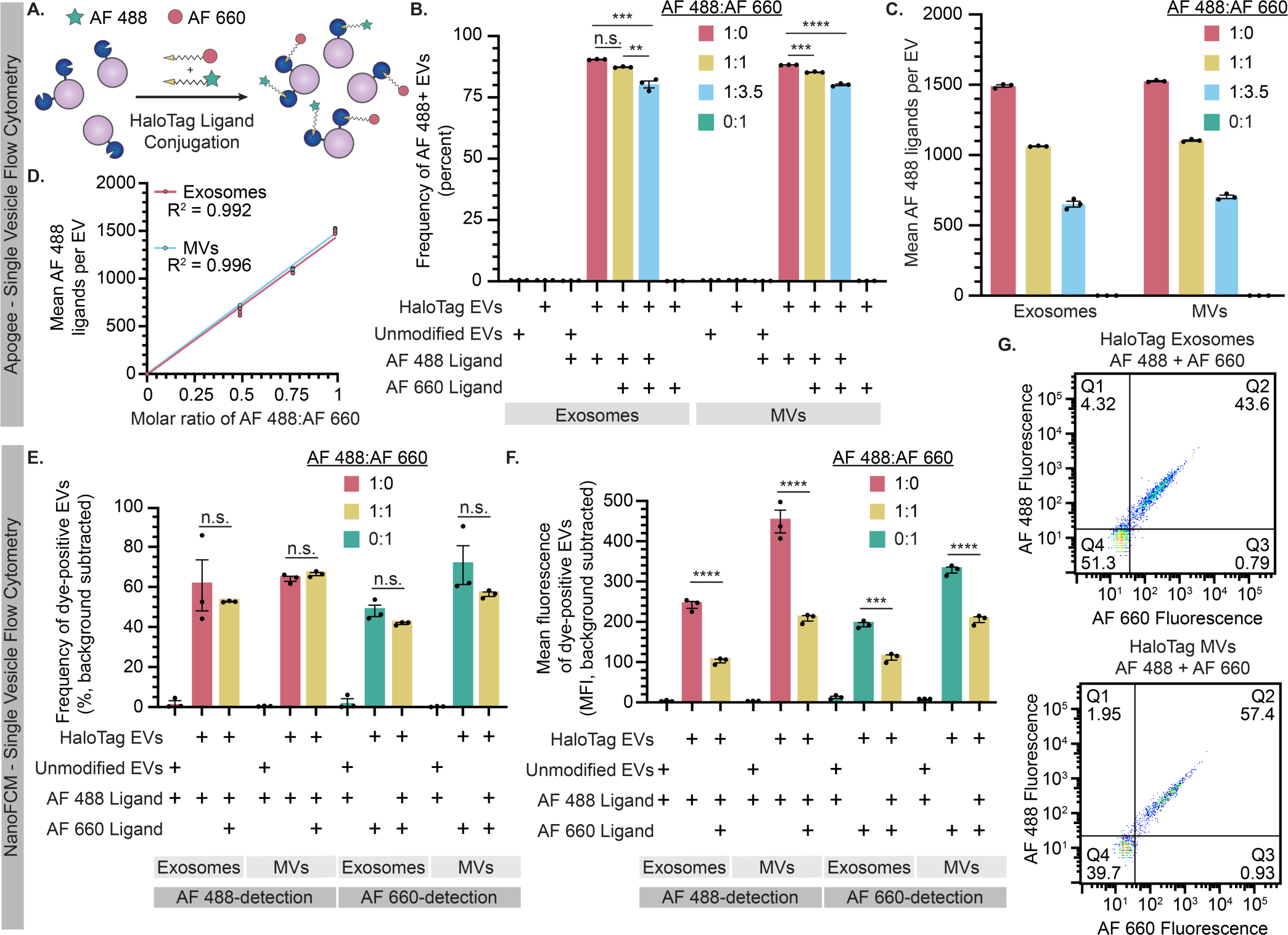
HaloTag EVs can be functionalized with multiple ligands. A) Cartoon illustrating the hypothesis that tuning the stoichiometric ratio of different ligands in a reaction may control relative ligand conjugation onto HaloTag EVs. B-D) Analysis of EVs by Apogee SV-FC. B) Frequency of AF 488+ EVs minimally decreases upon increasing AF 660 ligand concentration, which is consistent with overall high levels of HaloTag display and conjugation. C) Increasing the relative amount of AF 660 ligand proportionally decreases the average amount of AF 488 conjugated to the HaloTag EVs with tradeoffs proportional to their known relative rates of reaction. D) AF 488 conjugation scales linearly with the fraction of AF 488 ligand (vs. competitor AF 660 ligand) in the reaction. E-F) Analysis of EVs by NanoFCM SV-FC including frequency (E) and efficiency (F) of dye labeling evaluated at the single EV level. Note that the magnitudes of fluorescence detected in each channel (AF 488 vs. AF 660) cannot be compared across channels (i.e., the fluorescence units are independently scaled based upon instrument settings). G) HaloTag EVs can be co-labeled by AF 488+ and AF 660+ HaloTag ligands. EV experiments were performed in technical triplicate. Error bars indicate standard error of the mean. Pairwise comparisons were made using unpaired Student’s t-test. Multicomparison statistical analysis was performed using a one-way ANOVA test, followed by Tukey’s multiple comparison test to evaluate specific comparisons (*p < 0.05, **p < 0.01, ***p < 0.001, ****p < 0.0001).

We next sought to confirm co-labeling of EVs by fluorescence-based detection of both ligands. To evaluate co-labeling of a population of EVs, HaloTag EVs were conjugated to fluorescent dyes, and equal counts of EVs were run on an SDS-PAGE gel, after which fluorescence of the corresponding bands were quantified by imaging using a previously established method.^20^ Both AF 488 and AF 660 were detectable in co-labeled EV samples, and single-labeled EV samples were fluorescent only in the corresponding channel (**Figure S13A**). Absolute levels of EV labeling (at the population or bulk level) were estimated using a calibration curve based on conjugated HaloTag standard protein (**Figure S13B**). This method yielded somewhat lower estimates of average of HaloTag proteins per EV compared to SV-FC analysis, and this difference was most pronounced for MV samples (**Figure S13C**). In agreement with SV-FC observations, we observed a proportional decrease in the average number AF 488 per EV when the AF 660 ligand was included, supporting our prior interpretation of co-labeling experiments. Interestingly, AF 660-conjugation resulted in substantially lower estimates of ligands per EV compared to the same analysis using an AF 488 ligand. Since our samples fell within the ranges covered by the relevant standard curves, it is unlikely that detection sensitivity can explain these discrepancies. Since the ligands appeared to compete for HaloTag sites on EVs as expected (as indicated by decrease in AF 488 signal when co-labeling was performed), a possible explanation is that the AF 660 ligand competes for active HaloTag sites on EVs in manner that does not necessarily result in a complete conjugation reaction; this could be consistent with the manufacturer’s report that this ligand exhibits lower rates of reaction compared the AF 488 ligand. Altogether, these analyses indicate that co-labeling of EVs is possible at the bulk level—co-labeling of any single EV is not yet confirmed by this assay—and it provides insights as to how quantification should be interpreted as a function of ligand choice and labeling protocol. To evaluate co-labeling of HaloTag EVs at the single vesicle level, we next employed a NanoFCM Flow Nanoanalyzer. This SV-FC instrument is capable of detecting two channels of fluorescence—both AF 488 and AF660 dyes, in this case. Compared to the Apogee SV-FC analysis, we observed a somewhat lower frequency of HaloTag+ EVs in single dye-labeled samples (∼50-70%) (**Figure 3E**), but the degree of labeling (i.e., magnitude of single color fluorescence) again decreased upon co-labeling with both ligands, reflecting competition for limited HaloTag sites (**Figure 3F**). One possible explanation for the difference in overall labeling efficiency is that the NanoFCM can better identify smaller particles in our samples, which may be a distinct type of particle or contaminating debris. Most notably, we directly confirmed the presence of co-labeled EVs; the majority of EVs were both AF 488+ and AF 660+ (**Figure 3G**). We could not quantify the absolute amount of AF 488 labels per EV on the NanoFCM, since suitable quantification beads are not available for this instrument (which has a smaller maximum particle size than does the Apogee). Overall, these results confirm that individual HaloTag EVs can be co-labeled and corroborate and complement results observed using the Apogee SV-FC.

We next compared our results to those one might obtain with a conventional flow cytometer, using an instrument configured to detect EVs.^50^ A key challenge in such approaches is that compared to cells, EVs scatter light to a profoundly lower extent, and thus distinguishing EVs from bubbles and debris using scattering is challenging.^28^ Although methods such as lipophilic dye labeling of EVs provide an alternative to scattering-based identification of EVs, we opted to avoid potential spectral overlap challenges for this assay and instead employed methods comprising stringent sheath fluid filtering and a gating strategy focused on the specific goal of evaluating co-labeling (**Figure S14A**). SSC-A size gating was employed to minimize non-EV events while minimizing EV event exclusion, resulting in a population of high-scattering EVs within our size range of interest (**Figure S9A**). Approximately 90% of high-scattering HaloTag exosomes were labeled with AF 488, while a lower labeling frequency of ∼60% was observed for high-scattering MVs (**Figure S14B**). As observed via Apogee SV-FC, the frequency of AF 488+ EVs decreased when these EVs were co-labeled with AF 660 (**Figure S14B**). We also observed co-labeling of some EVs with both AF 488 and AF 660 ligands, but to a lower degree of co-labeling than observed via NanoFCM (**Figure S14C**), which is possibly attributable to limits of detection associated with characterizing EVs on this instrument. Overall, this comparative analysis across several EV characterization platforms demonstrates that the HaloTag system enables titration of multiple ligands for multi-functionalization of the EV surface.

### Antibody labeling of EVs underestimates EV display

Having shown that HaloTag EVs can be used as a modular display platform, we harnessed this technology to evaluate a key question in EV quantification—how does quantification of EV surface display by antibody labeling compare to the true level of display? Fluorescently labeled antibodies enable labeling endogenous and engineered proteins on the surface of cells for analysis by flow cytometry, and this method can be made quantitative using calibration standards. This overall approach can be extended to EVs for analysis by SV-FC.^48,51–53^ Given the large difference in diameter between mammalian cells and EVs (10-100 μm depending on cell type vs ∼100 nm for EVs), and the relatively large size of antibodies (∼10 nm for IgG), it is critical to evaluate the extent to which antibodies can be used to probe EVs at a single vesicle level.^54,55^ To investigate this question, an AF 488-conjugated anti-FLAG antibody was used to target the 3x FLAG tag on the N-terminus of the HaloTag construct (**Figure S15A**). For HaloTag-expressing cells, fluorescence attributed to labeling was 10-fold greater for cells labeled with HaloTag ligand compared to cells labeled with antibody. Background labeling of unmodified HEK293FT cells was also 8.5-fold lower when labeled with HaloTag ligand vs antibody (**Figure S15B**). Thus, compared to antibody labeling of cells, HaloTag labeling yields both higher efficiency and lower background.

Similar trends were observed for antibody labeling of HaloTag EVs detected via Apogee SV-FC. Antibody-mediated labeling yielded substantially lower labeling for both exosomes and MVs (**Figure S15C-D**). The non-specific antibody binding to unmodified EVs was also higher than that for HaloTag ligand for both exosomes and MVs (**Figure S15D**). Notably, these trends hold even though each antibody is conjugated to multiple fluorophores (this antibody lot contains 6.24 moles of fluorophore per mole of antibody). After correcting for this fact, we can estimate the number of labeling events detected by each method (**Figure S15E**).^56^ Overall, antibody-mediated labeling of HaloTag EVs underestimates the level of HaloTag display (compared to that determined by HaloTag ligand labeling) in terms of both frequency of EVs that are HaloTag+ (underestimates by ∼57% of the EV population) and the level of HaloTag display (underestimates by ∼800-850 labeling events per vesicle).

Given the substantial non-specific binding observed with the anti-FLAG antibody, we investigated whether probing using other common antibodies also underestimates EV display of HaloTag. To this end, an HA tag was cloned in place of the N-terminal FLAG tag on the HaloTag construct, which was then expressed by transient transfection of HEK293FT cells to produce EVs loaded with HA tag-HaloTag proteins (**Figure 4A**). HA tag-HaloTag display was detectable on transfected cells using either an anti-HA antibody or HaloTag ligand (**Figure 4B**). When analyzed by NanoFCM SV-FC, anti-HA EV labeling yielded substantially higher background for both exosomes and MVs compared to HaloTag ligand labeling (**Figure 4C**). As observed with anti-FLAG labeling, anti-HA antibody labeling of HaloTag EVs underestimated the frequency of HaloTag display by ∼50% compared to HaloTag ligand labeling (**Figure 4D**). By factoring in the average fluorophores per antibody (3.53 fluorophores per antibody for this antibody lot), we calculated that anti-HA antibody labeling under-estimates the average magnitude of HaloTag proteins displayed per EV for both exosomes and MVs (by 2.3 and 1.5-fold, respectively) (**Figure 4E**). Altogether, the anti-HA antibody outperformed the anti-FLAG antibody, due at least in part to reduced non-specific binding, but it still underestimated both the frequency of HaloTag+ EVs and the efficiency of HaloTag protein display per EV. Although we expect that such differences might be most pronounced for a highly expressed protein such as our HaloTag system, and it is possible that careful antibody titration could further reduce these underestimates on a case-by-case basis, these observations generally provide important context for interpreting experiments employing antibody-based labeling of EVs.

**Figure 4:**
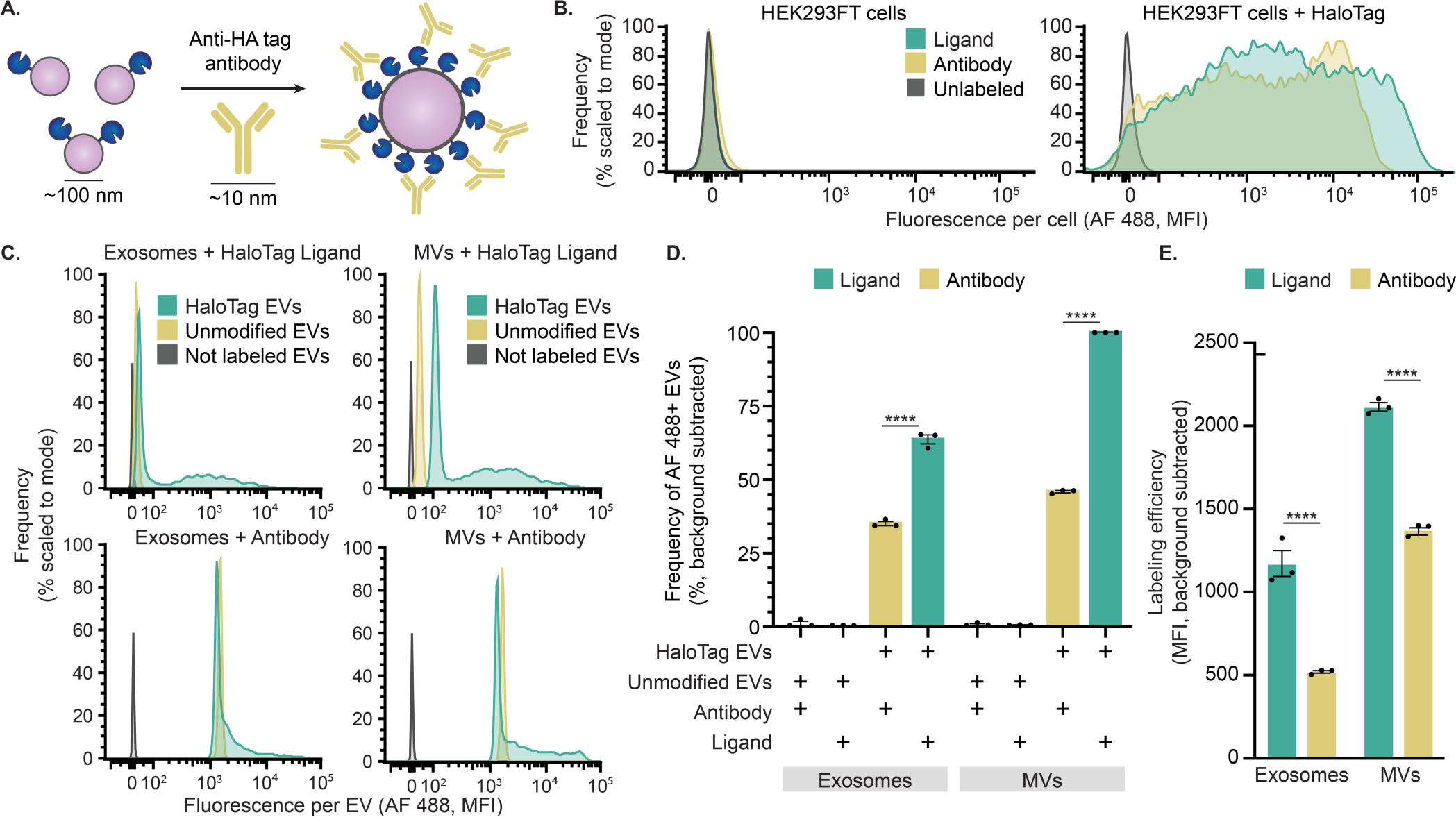
Antibody labeling of HaloTag EVs underestimates protein display. A) This cartoon illustrates anti-HA tag antibody-mediated labeling of HaloTag protein on EVs. HaloTag protein (∼4 nm) and EVs (∼100 nm) are not to scale. B) Antibody labeling of EV producer cells compared to HaloTag conjugation. C) Antibody labeling of EVs compared to HaloTag conjugation; all analyses employ NanoFCM flow cytometer. D) Antibody labeling underestimates the frequency of HaloTag+ EVs relative to HaloTag-conjugation. E) Antibody labeling underestimates efficiency of HaloTag display; labeling efficiency was calculated by scaling the fluorescence (MFI) by the fluorophores per label (1 for HaloTag ligands; 3.53 for antibody-labeling given characterization of this antibody lot). Cell experiments were performed in biological triplicate. EV experiments were performed in technical triplicate. Error bars indicate standard error of the mean. Pairwise comparisons were made using unpaired Student’s t-test. Multicomparison statistical analysis was performed using a one-way ANOVA test, followed by Tukey’s multiple comparison test to evaluate specific comparisons (*p < 0.05, **p < 0.01, ***p < 0.001, ****p < 0.0001).

### Microscopy methods corroborate HaloTag-based characterization of EVs

To validate our SV-FC results, we next utilized orthogonal microscopy methods to quantify the number of HaloTag proteins per EV at a single vesicle resolution. Fluorescence microscopy is a technique used to measure the fluorescence of dye-labeled EVs and quantify the number of attached fluorophores.^57–59^ Using this approach, we imaged individual AF 488 labeled HaloTag EVs, and fluorescence was quantified using the same beads used for SV-FC. Fluorescence microscopy yielded similar lognormal distributions of EV functionalization compared to those observed by SV-FC (**Figure 5A**). In addition, the average numbers of HaloTag proteins per EV as measured by fluorescence microscopy were comparable to values observed by SV-FC, for both EV subpopulations (**Figure 5B**).

**Figure 5:**
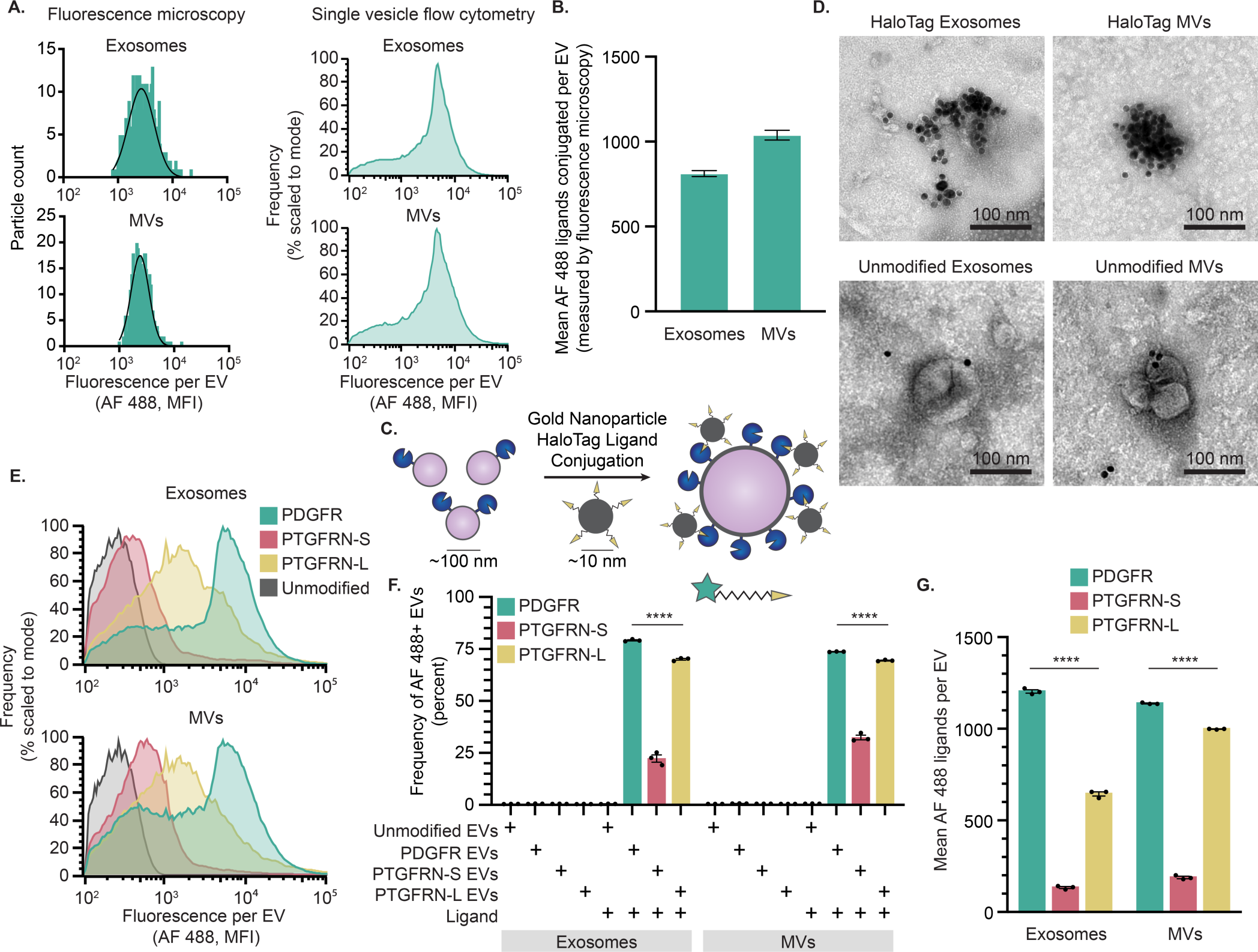
HaloTag-mediated quantification of EV surface display can be corroborated and employed to study EV engineering design choices. A) Distributions of fluorescence attributable to HaloTag conjugation as measured by fluorescence microscopy are comparable to those obtained by SV-FC for both exosomes and MVs. A minimum of 300 events were analyzed by fluorescence microscopy. B) Quantification of HaloTag copies per EV determined via fluorescence microscopy is comparable to that determined by SV-FC. C) This cartoon illustrates AuNP HaloTag ligand conjugation to HaloTag EVs. HaloTag protein (∼4 nm) and EVs (∼100 nm) are not to scale. D) EVs conjugated to gold nanoparticle HaloTag ligands show surface functionalization for HaloTag EVs (top) and minimal association with unmodified EVs (bottom) by TEM. Scale bar: 100 nm. E) PTGFRN-S TMD and PTGFRN-L TMD HaloTag display constructs yield lower levels of HaloTag display compared to PDGFR TMD HaloTag construct, as measured by SV-FC. F) The PTGFRN-S TMD display construct yields fewer HaloTag+ EVs compared to the PDGFR TMD and PTGFRN-L TMD constructs. G) PTGFRN-L display loads HaloTag protein onto MVs moreso than exosomes, whereas PTGFRN-S loads HaloTag comparably into both EV populations, as does PDGFR display (albeit at higher levels). All experiments were performed in technical triplicate. Error bars indicate standard error of the mean. Multicomparison statistical analysis was performed using a one-way ANOVA test, followed by Tukey’s multiple comparison test to evaluate specific comparisons (*p < 0.05, **p < 0.01, ***p < 0.001, ****p < 0.0001).

To further test our interpretation that HaloTag labeling leads to EV surface functionalization, we next evaluated labeling using a scheme that can be visualized by TEM, as this approach is able to distinguish EVs from non-EV materials that could co-precipitate during purification.^60–62^ Immunogold labeling of EVs enables visualization of proteins on the surface of EVs and their relative abundance.^63^ Since no commercial immunogold HaloTag ligand exists, we synthesized a custom gold nanoparticle (Au-NP) functionalized with a HaloTag ligand for immunogold labeling using established methods (**Figure 5C**).^64,65^ Highly Au-NP functionalized HaloTag EVs were observed via TEM after conjugation, while unmodified EV samples exhibited minimal co-localization between EVs and Au-NPs (**Figure 5D**).

Although it was not possible to perform a systematic, unbiased quantification using this TEM approach, we did note that these clusters generally contain fewer Au-NP per EV than would be expected from the AF 488 SC-FV analyses, indicating that Au-NP labeling provides qualitative confirmation of general phenomena but may not be suitable for quantification of EV display. Overall, confirmation of the SV-FC results by orthogonal microscopy methods shows that HaloTag-based labeling is robust across multiple EV analysis methods.

### Employing HaloTag to evaluate EV display design choices

When displaying proteins on the surface of EVs, design choices can influence outcomes such as level of protein display and distribution of display across EV subsets. Choices such as transmembrane domain (TMD) used, linker used to fuse the cargo and TMD, and cytoplasmic residues included can all modulate protein loading and display on EVs.^66,67^ Having validated the HaloTag system as a method for characterizing EVs, we next employed this system to quantitatively evaluate the relationship between common design choices and EV surface display. As a test case, we compared the PDGFR TMD-based display system to one based upon Prostaglandin F2 receptor negative regulator (PTGFRN), another EV-associated protein that has been reported to be highly efficient for displaying cargo on EVs.^14^ Previous reports analyzed PTGFRN truncation mutants as display candidates^14^ but did not investigate a minimal PTGFRN TMD-based display scaffold, so to explore this possibility, entry (Q9P2B2) in UniProt was analyzed using several protein prediction software packages (UniProt, TMdock, HMMtop) to predict the TMD region. This analysis yielded two possibilities—a shorter domain prediction (termed, PTGFRN-S) and a longer, more conservative prediction (termed, PTGFRN-L). These TMDs were cloned into the HaloTag construct, after which stable cell lines and EVs were produced as with PDGFR construct. For HaloTag-expressing cells conjugated to ligand, the PDGFR construct resulted in a 20.4-fold larger fluorescence intensity compared to the PTGFRN-S construct and 1.4-fold for the PTGFRN-L construct (**Figure S16**). The engineered HaloTag constructs were detected in all cell lysates at comparable levels via Western blot, but the engineered protein was enriched in the PTGFRN-S EVs compared to the PDGFR and PTGFRN-L EVs (**Figure S17**). This analysis confirmed that all constructs appear nominally functional, and so all were carried forward to evaluate EV surface display.

We evaluated the consequences of TMD choice on protein display at the single EV level using Apogee SV-FC. Compared to the PDGFR HaloTag construct, we observed a significant decrease in percent of fluorescent HaloTag+ EVs for the PTGFRN-S EVs (22% of exosomes and 32% of MVs were HaloTag+), but a relatively mild decrease in labeling frequency for PTGFRN-L EVs, (70% of exosomes and 69% of MVs were HaloTag+) (**Figure 5E-F**), agreeing with the trends observed in cells. A significant decrease was also observed for the extent of HaloTag protein loading of PTGFRN-S EVs, which contained approximately 10 times less HaloTag proteins/EV as PDGFR EVs for exosomes and 6 times less for MVs. Interestingly, the PTGFRN-L TMD conferred different amounts of HaloTag display between the EV populations, with MVs displaying ∼350 more HaloTag proteins per vesicle compared to exosomes in a pool of EVs produced from the same cells (**Figure 5G**). All together, these results indicate that the additional amino acids in the PTGFRN-L TMD confer high construct loading onto EVs compared to the PTGFRN-S sequence, and that PTGFRN-L loads protein comparably to PDGFR into MVs but not exosomes. These precise and unambiguous comparisons highlight the quantitative characterization capabilities of the HaloTag system and its utility in guiding future EV display system design choices.

## DISCUSSION

HaloTag display on EVs combined with single vesicle analysis and quantification enables one to investigate EV population heterogeneity and protein loading in a manner that complements classically used EV analysis techniques. The high frequencies of EVs displaying engineered HaloTag protein (∼90% HaloTag+ for both exosomes and MVs) observed here are consistent with previously reported measurements based upon SV-FC combined with other labeling methods, with highly efficient display platforms leading to 95-98% of EVs loaded with fluorescent protein fused to EV membrane proteins.^14,27^ Previously reported bulk EV measurements (e.g., based upon ELISA) reported similar average levels of protein loading onto EVs, with the most highly efficient display systems (including full length PTGFRN or a distinct Δ687 truncation mutant) loading 600-1000 GFP fusion proteins per EV.^14^ Overall, our single vesicle measurements for HaloTag loading onto EVs agree with previously observed highly-efficient display scaffolds, but the HaloTag system allows us to make further quantitative observations about the distribution of protein loading levels across the EV population.

Although antibodies are useful in many EV labeling applications, antibody-mediated quantification of abundant surface proteins on EVs likely underestimates surface display. We hypothesize that labeling by reversible antibody binding is limited by equilibrium (i.e., local surface concentration becomes very high as labeling increases) and/or steric hindrance given the relative sizes of antibodies and EVs. Affinity reagents such as nanobodies, which are ∼10x smaller than antibodies and retain high target specificity, could potentially bypass some of the steric limitations of traditional antibodies.^28,68^ It is possible that one could use model systems similar to the one reported here, in which a HaloTag is fused to a target epitope on the EV surface, to directly compare expressions levels evaluated by HaloTag ligand conjugation versus affinity reagent-mediated labeling as a means of calibrating such a labeling scheme. Such analyses align with growing interest in assays enabling single EV characterization and absolute quantification.

Combining bulk analyses and single EV analyses provides insights exceeding those provided by either approach alone. For example, bulk analysis (by western blot) indicated that the PTGFRN-S scaffold yielded high HaloTag display on (or in) EVs, yet comparisons by SV-FC suggested that much of the HaloTag fused to this scaffold is non-reactive with cell-impermeable ligand. It is possible that the shorter TMD used in this construct (which is shorter compared to the PTGFRN-L scaffold, for example) causes HaloTag to be incorporated in an orientation or form that precludes reaction with ligand. In general, it is possible that the location of the HaloTag protein in the fusion construct or the size of the HaloTag (∼33 kDa) could impact translation, folding, and loading of this domain on EVs (i.e., compared to display of other proteins using the same system). Although we aimed to mitigate any inhibitory effects on display by placing the HaloTag protein at the N-terminus and using a helical linker to facilitate proper folding, display of other domains could certainly vary based upon similar considerations. These comparisons also yielded novel observations that could drive further investigation to understand the impact of specific residues on protein loading onto EVs. For example, PTGFRN-L loads HaloTag at comparable levels into MVs as does PDGFR does, but the former scaffold loads this tag at significantly lower levels for exosomes. The use of HaloTag EVs to investigate the effects of display platform design choices on cargo loading and functionality may increase understanding of EV biology and biogenesis, and such insights for engineering EV surface-display can yield improved biotechnologies.

In addition to the analytical utility described in this study, HaloTag EVs could also provide a simple yet highly specific system for functionalization of EVs with various surface moieties of interest, ranging from peptides to proteins to synthetic molecules. The ability to generate many different populations of functionalized EVs from one parental HaloTag pool by simply changing the HaloTag ligand is appealing, as this would circumvent the need to repeat the design process for each unique construct one wishes to display on EVs and would reduce the number of EV harvests required, both of which can be time consuming and costly. Conjugation of cargo onto HaloTag EVs after biogenesis also avoids several biomanufacturing challenges. For example, large proteins and proteins which are toxic to cells can be difficult to express and package on EVs during biogenesis, resulting in low levels of protein loading onto EVs or cell death leading to low EV yields. HaloTag EV surface conjugation of a recombinant protein of interest could reduce these challenges. Further, HaloTag-mediated display on EVs facilitates tunable loading of different molecules onto the same EV (leveraging many copies of HaloTag/EV and irreversible conjugation). This approach could be used to generate EVs with defined ratios of displayed molecules, which could meet some application-specific needs that would otherwise be challenging to implement with genetic engineering alone. HaloTag-mediated functionalization of the EV surface provides a modular system for engineering EVs and can benefit both fundamental research and the development of EV therapeutics.

## ACKNOWLEDGEMENTS

This work was supported in part by a Cornew Innovation Awards from the Chemistry of Life Processes Institute (Northwestern University). R.E.M. was supported in part by the Northwestern University Graduate School Cluster in Biotechnology, which is affiliated with the Biotechnology Training Program; by the National Institutes of Health Training Grant (T32GM008449) through Northwestern University’s Biotechnology Training Program; and by a Ryan Fellowship from the International Institute for Nanotechnology at Northwestern University. D.M.S. was supported by the National Science Foundation Graduate Research Fellowship under Grant No. (D.M.S. # DGE-1324585). B.N.D. was supported by the National Science Foundation Graduate Research Fellowship under Grant No. (B.N.D. # DGE-2234667). This work was supported by the Northwestern University Flow Cytometry Core Facility supported by Cancer Center Support Grant (NCI CA060553). This work made use of the BioCryo facility of Northwestern University’s NUANCE Center, which has received support from the SHyNE Resource (NSF ECCS-2025633), the IIN, and Northwestern’s MRSEC program (NSF DMR-1720139). Biological and chemical analysis was performed in the Analytical bioNanoTechnology Core Facility of the Simpson Querrey Institute at Northwestern University. The U.S. Army Research Office, the U.S. Army Medical Research and Materiel Command, and Northwestern University provided funding to develop this facility and ongoing support is being received from the Soft and Hybrid Nanotechnology Experimental (SHyNE) Resource (NSF ECCS-1542205). This work was supported by the Northwestern University Sanger Sequencing Facility. This work was supported by the Northwestern University Keck Biophysics Facility funded by the NIHS10 OD026963 Grant. We acknowledge support from the National Institutes of Health Grant (National Institute of Neurological Disorders and Stroke) 5R01NS115571. This work made use of the Center for Synthetic Biology BioFoundry facility at Northwestern University, which has received support from the Army Contracting Command (W52P1J-21-9-3023). Any opinion, findings, and conclusions or recommendations expressed in this material are those of the authors and do not necessarily reflect the views of the National Science Foundation. The authors kindly thank the Leonard lab members and Dr. Suchitra Swaminathan for useful discussions throughout the planning, experimental, analysis, and writing phases of this project.

## DATA AVAILABILITY STATEMENT

The data that support the findings of this study will be made openly available at Zenodo and will be published along with the final version of this manuscript. The plasmids generated in this study will be deposited with and distributed by Addgene at the time of publication, including complete and annotated sequence files, at https://www.addgene.org/Joshua_Leonard/.

