## Supplementary Information for "HaloTag display enables quantitative single-particle characterization and functionalization of engineered extracellular vesicles"

**Supplementary Information**
**For**

### HaloTag display enables quantitative single-particle characterization and functionalization of engineered extracellular vesicles

Roxana E. Mitrut[1,2], Devin M. Stranford[1,2,†], Beth N. DiBiase[1,2,†], Jonathan M. Chan[1], Matthew D. Bailey[3], Minrui Luo[3,5], Clare S. Harper[4], Thomas J. Meade[3,4,5,6], Muzhou Wang[1], Joshua N. Leonard[1,2,4,5,6]*

1. Department of Chemical and Biological Engineering, Northwestern University, Evanston, IL, 60208, USA
2. Center for Synthetic Biology, Northwestern University, Evanston, IL, 60208, USA
3. Department of Chemistry, Northwestern University, Evanston, IL, 60208, USA
4. Interdisciplinary Biological Sciences Training Program, Northwestern University, Evanston, IL, 60208, USA
5. Chemistry of Life Processes Institute, Northwestern University, Evanston, IL, 60208, USA
6. Member, Robert H. Lurie Comprehensive Cancer Center, Northwestern University, Evanston, IL, 60208, USA

†These authors contributed equally as co-second authors.

Contents

**Supplementary Methods**

**Supplemental Figure 1**

**Supplementary Scheme 1**

**Supplemental Figure 2**

**Supplementary Scheme 2**

**Supplemental Figure 3-4**

**Supplementary Figures 5-17**

#### SUPPLEMENTARY METHODS

##### General Methods

Unless otherwise noted, reagents used were commercially purchased at reagent grade or higher and used without further purification. All water used was type 1 ultrapure water (≥ 18.2 MΩ) from a Synergy® UV Water Purification System (Millipore-Sigma). Chemical intermediates were purified by flash chromatography using silica gel 60 (0.04-0.063 mm, Macherey-Nagel GmbH & Co KG) by the methods from the corresponding references. NMR spectra were collected using a Bruker Avance III 500 MHz instrument equipped with a DCH CryoProbe. NMR samples were prepared in deuterated chloroform ($CDCl_3$) containing an 0.03% tetramethylsilane (v/v). Chemical shifts are reported in reference to residual solvent signal ($CDCl_3$ $^1H$: 7.26, $^{13}C$: 77.0). NMR splitting was assumed to be first order and apparent multiplicity is reported as "brs" = broad singlet, "t" = triplet, "m" = multiplet. UV/visible spectroscopy was collected on an Agilent Cary 60 UV-Vis Spectrophotometer using a microdrop cell (TrayCell 105.800-UVS, Hëllma Analytics) on volumes of 4 µL at a path length of 1 mm. TEM micrographs were collected on a Hitachi HD-2300 Dual EDS Cryo STEM. TEM image workup for particle size was accomplished via ImageJ software.[1] Centrifugation was carried out using an Eppendorf 5810 R Centrifuge. Liquid rotations were performed with a Fisherbrand™ Multi-Purpose Tube Rotator (~5 rpm). ESI-MS was collected on a Bruker AmaZon SL instrument in methanol. Abbreviations: gold nanoparticles (AuNPs), *tert*-butyloxycarbonyl substituent group (Boc), di-*tert*-butyl dicarbonate ($Boc_2O$), fluorenylmethoxycarbonyl substituent group (Fmoc), *N*,*N*-dimethylformamide (DMF), tetrahydrofuran (THF), 1X phosphate-buffered saline (PBS), trifluoroacetic acid (TFA), triisopropylsilane (TIPS), triethylsilane (TES), surface plasmon resonance (SPR), diameter (d), electrospray ionization mass spectrometry (ESI-MS), thin-layer chromatography (TLC), calculated (calcd), parts per million (ppm), equivalents (eq.), circa (*ca.*).

##### Synthesis of Citrate-Stabilized AuNPs

Gold nanoparticles (AuNPs) were synthesized by a variation of the Frens/Turkevich method.[2] All used glassware was pre-washed with aqua regia and triple rinsed with water before synthesis. $HAuCl_4$ hydrate (42.2 mg, 0.11 mmol) was dissolved in water (99 mL). The clear, yellow solution was brought to a rolling boil with stirring and allowed to boil for 15 min. To the boiling solution a solution of tribasic sodium citrate dihydrate (115.3 mg, 0.43 mmol) in water (1 mL) was added quickly. Addition of the citrate solution was followed by the typical loss of solution color followed by a greying of solution until it turned black and ultimately took on a wine-red color. The solution was allowed to continue to reflux for 15 minutes after citrate solution addition. The wine-red solution was then allowed to slowly cool to room temperature and filtered through a medium glass frit to obtain the colloid.

##### Preparation of TEM Samples

To prepare the samples for transmission electron microscopy (TEM) microscopy, carbon-coated, copper TEM grids were used as a support film. However, these grids are naturally hydrophobic, making it difficult for aqueous solutions of particles to uniformly spread and adhere to the grid surface. Therefore, grids were glow discharged on a Pelco EasiGlow Glow Discharge system for 30 seconds to deposit a charge onto the grids, rendering them hydrophilic. Next, 5 µL of the AuNP solution was placed onto the grid, and excess solution was absorbed by Whatman filter paper. The sample was then left to settle for 1-2 minutes to dry. TEM micrographs of the synthesized AuNPs were collected using a Hitachi HD-2300 Dual EDS Cryo STEM operated at a voltage of 200kV.

#### Characterization of Citrate-Stabilized AuNPs

Citrate-stabilized AuNPs were characterized by TEM for gold core size (**Figure S1A-B**). Raw micrographs were worked up with ImageJ software.[1] Scale bars were set and the images were then converted to 8-bit. A gaussian blur of 2 was applied and the image threshold was adjusted to create a binary image of black particles on a white background. Binary images were then converted to a mask and the watershed function was applied to separate particles in close proximity. Particle areas were then collected from ImageJ, confirmed for overlap with raw images, and converted to diameters using the equation $d = 2 * \mathrm{sqrt}(\mathrm{Abs}/\boldsymbol{\pi})$ under the assumption of a circular projection of a spherical particle shape. The population of measured particles (N=133) was assumed to be gaussian and described with a mean of 12.3 ± 1.3 nm (error represents standard deviation around the mean). UV/visible spectroscopy was collected on the colloid to corroborate the average nanoparticle size as well as get a nanoparticle concentration via reported empirical equations (**Figure S1C-D**).[3] UV/vis was collected using a microdrop cell with a 1 mm path length. From the spectrum a calculated average size of 12.1 nm was determined in good agreement with the measured TEM value. From the UV/vis a calculated concentration of 23 nM AuNPs was determined using the reported extinction coefficient at 450 nm for 12 nm gold AuNPs.[3]

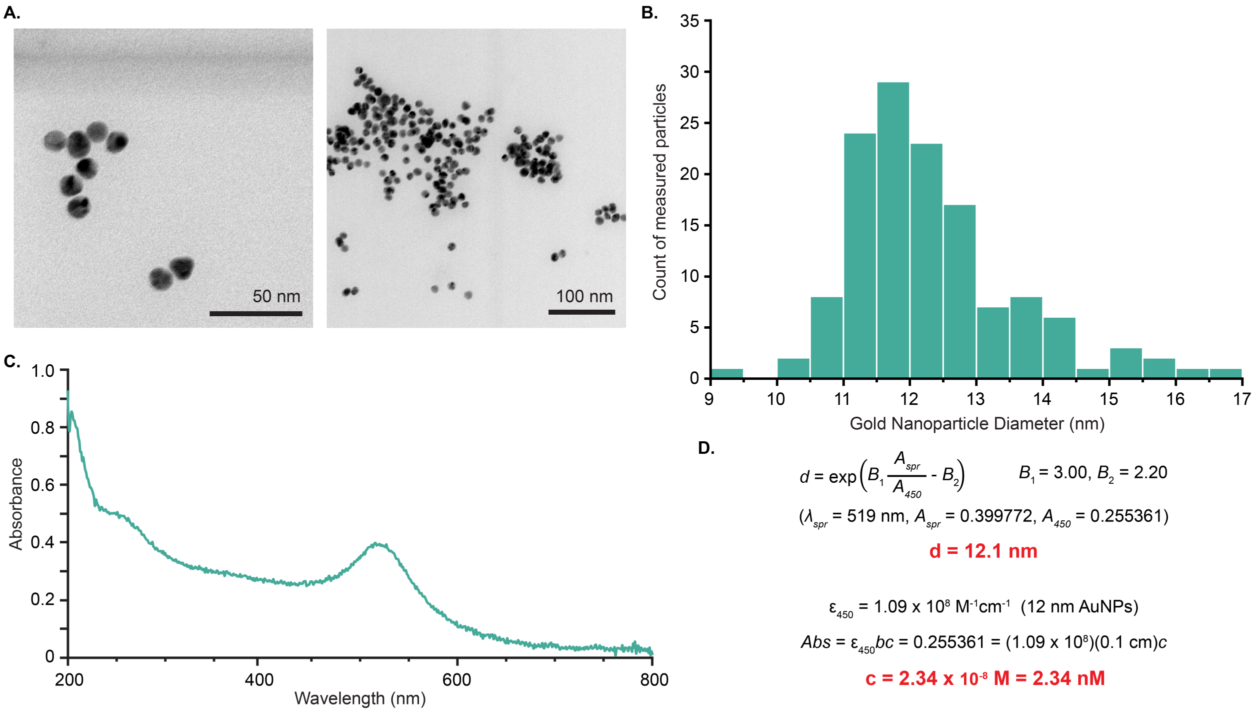


**Supplemental Figure 1:** Synthesized AuNPs display expected spherical morphology with a narrow size range. A) Raw TEM micrographs of the citrate-stabilized AuNPs. B) Histogram of measured AuNP core diameters (N=133). C) UV/visible spectrum of citrate-stabilized AuNPs in water, 1 mm path length. $\boldsymbol{\lambda}_{SPR}$ = 519 nm. D) UV/vis calculation of AuNP diameter and concentration.[3]

#### Synthesis of HT-ligand

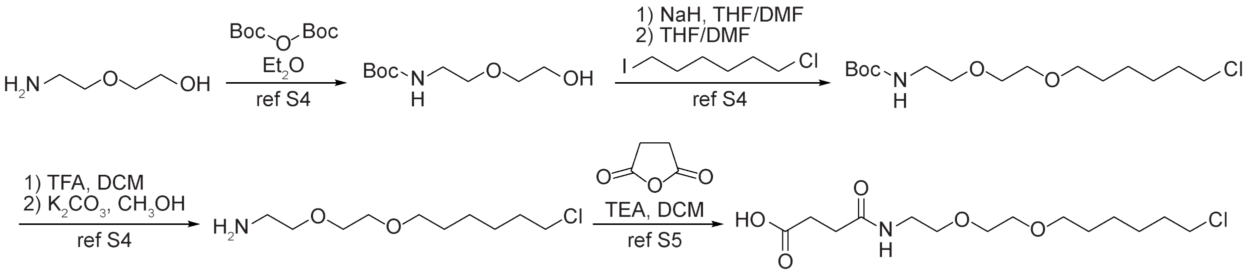


**Supplementary Scheme 1:** Synthetic route to coupling-ready ligand for HaloTag protein, **HT-ligand**.[4,5]

The synthesis of **HT-ligand** was accomplished in four steps from 2-(2-aminoethoxy)-ethanol following literature procedures (**Scheme S1**).[4,5] Each intermediate was isolated via standard phase silica gel chromatography by cited literature procedures. Briefly, the amine of 2-(2-aminoethoxy)-ethanol was protected using $Boc_2O$ in ethanol.[4] The unprotected alcohol was then deprotonated with sodium hydride and alkylated with 1-bromo-6-chlorohexane in a mixture of THF and DMF.[4] The Boc protecting group was then removed with 14% TFA in DCM (v/v) and then worked up with $K_2CO_3$ in methanol to yield a free amine.[4] The resulting free amine was further alkylated with succinic anhydride in dichloromethane in the presence of triethylamine to afford peptide coupling-ready **HT-ligand**.[5] The product was purified using flash chromatography on silica gel using 40:10:2 DCM/$CH_3OH$/$NH_4OH_{(aq)}$ and TLC was monitored using an eluent of 20:10:1 DCM/$CH_3OH$/$NH_4OH_{(aq)}$ and visualized with ninhydrin stain. $R_f$=0.6 (20:10:1 DCM/$CH_3OH$/$NH_4OH_{(aq)}$); $R_f$=0.2 (40:10:2 DCM/$CH_3OH$/$NH_4OH_{(aq)}$); $^1$H NMR (500 MHz, $CDCl_3$): δ=6.52 (brs, 1H), 3.65-3.57 (m, 4H), 3.57-3.41 (m, 8H), 2.67 (t, 2H, $^3J_{H,H}$=7.05Hz), 2.52 (t, 2H, $^3J_{H,H}$=7.10Hz) 1.82-1.73 (m, 2H), 1.67-1.57 (m, 2H), 1.50-1.41 (m, 2H), 1.41-1.32 (m, 2H); $^{13}$C NMR (125 MHz, $CDCl_3$) 175.1, 172.4, 71.4, 70.1, 70.1, 69.5, 45.0, 39.4, 32.5, 31.2, 30.4, 29.2, 26.6, 25.3.

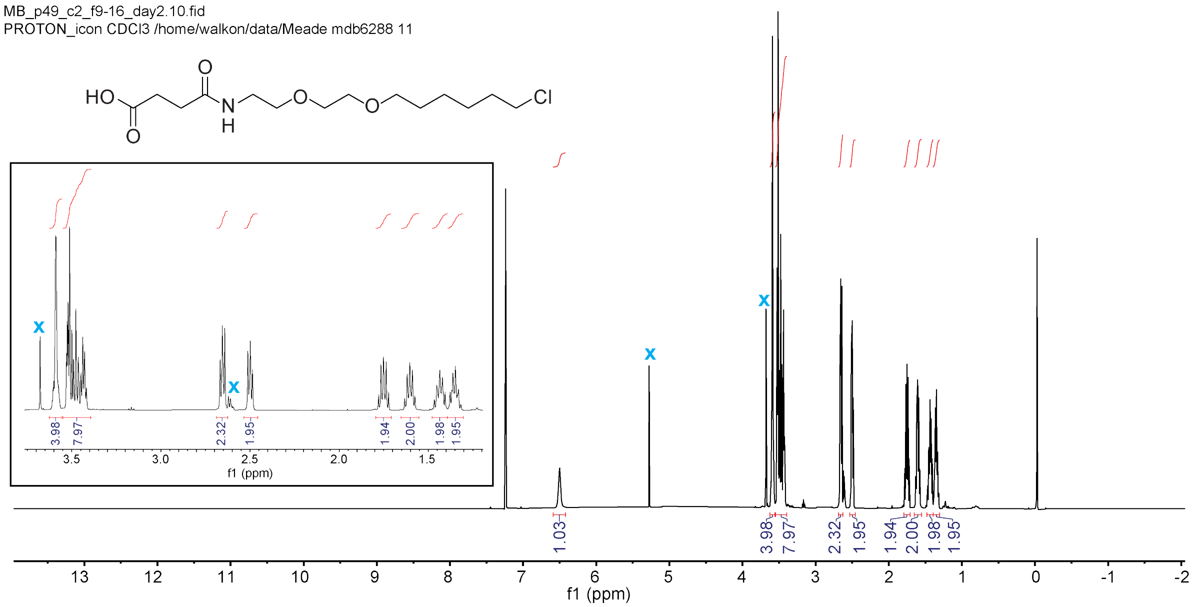


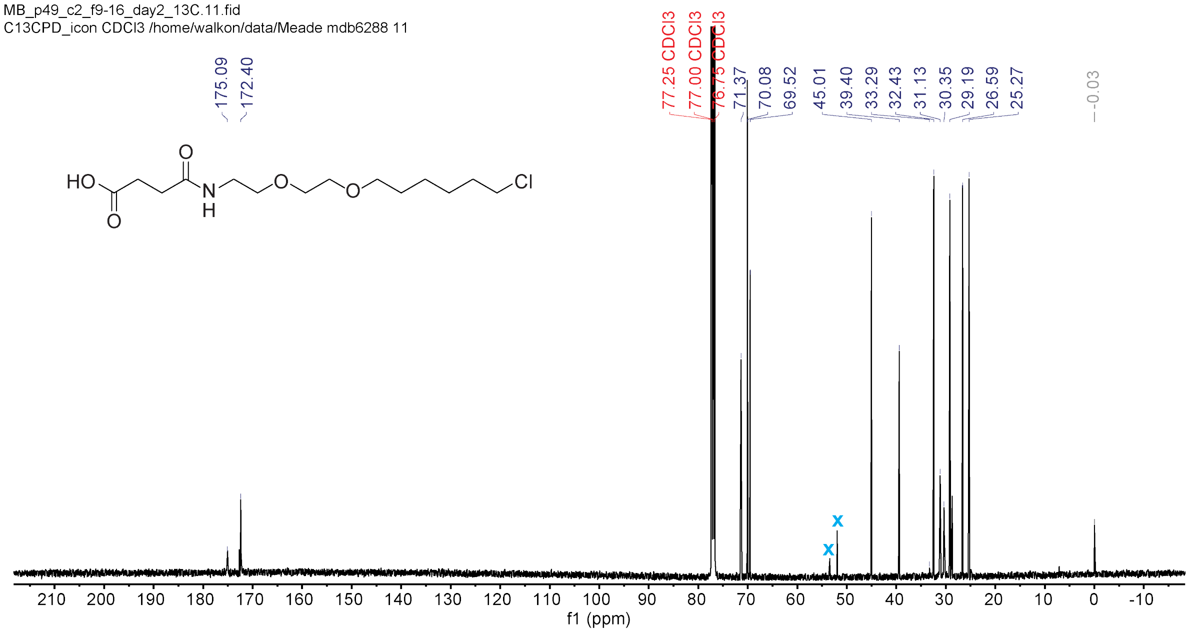


**Supplemental Figure 2:** $^{1}H$ NMR (top) and $^{13}C$ NMR spectra of **HT-ligand** in $CDCl_3$. The inset in the $^{1}H$ spectrum is a zoom-in of the peaks between 1.2-3.8 ppm. Blue Xs represent peaks which were ignored from residual DCM and an unknown minor impurity that co-eluted.

**Solid-Phase Peptide Synthesis**

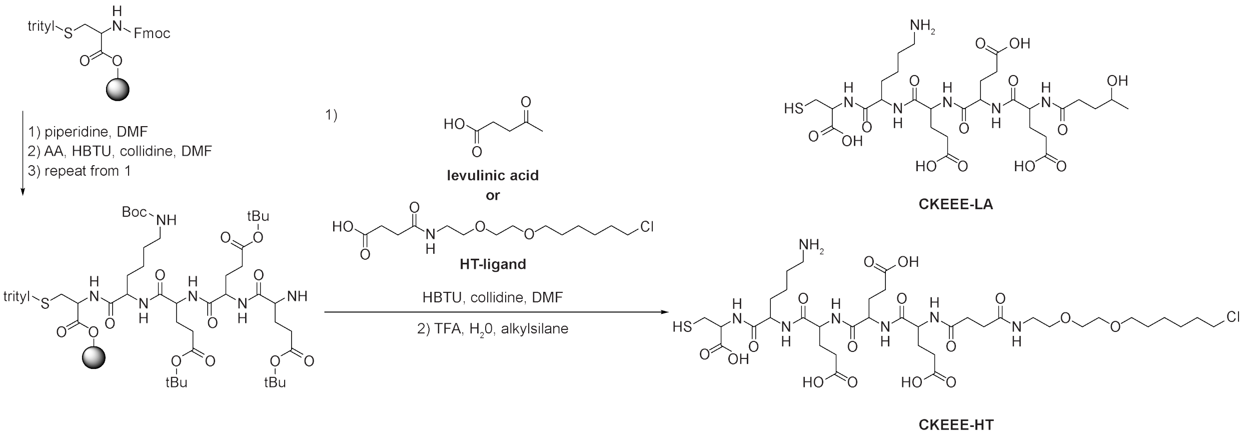


**Supplementary Scheme 2**. Reaction flow for solid-phase peptide synthesis for the **CKEEE-LA** and **CKEEE-HT** peptides**.**

Peptides were synthesized via solid-phase peptide synthesis (SPPS) using 2-chlorotritylchloride resin pre-loaded with S-trityl-L-cysteine (0.401 mmol/g, ChemImpex). Syntheses were carried out at a 0.1 mmol scale (*ca*. 250 mg of resin). The resin was swelled in a PolyPrep® chromatography column (BIO-RAD, 20 mL, 0.8×4.0 cm) in dichloromethane (15 mL) for 90 min and transferred to a glass peptide synthesis chamber affixed with a medium frit filter and valve selector between vacuum and high-pressure $N_2$ inlet. The resin was then washed with dichloromethane (3x10 mL) followed by anhydrous DMF (3x10 mL). Deprotections were carried out using a solution of piperidine in anhydrous DMF (10 mL, 20% v/v piperidine) with $N_2$ bubbling for 7 min. After deprotection the resin was washed five times with anhydrous DMF. Each washing involves the addition of 10 mL of anhydrous DMF, vigorously bubbling the mixture with $N_2$ gas, and removal of solvent with vacuum assistance. Amino acid couplings were performed with 4 eq. (0.4 mmol) of the relevant Fmoc-protected amino acid and 4 eq. (0.4 mmol) of HBTU in a solution of 2,4,6-collidine in anhydrous DMF (10 mL, 20% v/v 2,4,6-collidine, mixed fresh). Amino acid/HBTU solutions were allowed to sit for ≥15 min before use. Coupling solutions were added to the deprotected peptide and allowed to bubble with $N_2$ gas to mix for 20 min. After coupling, the same washing procedure was used as after deprotections. For each synthesis couplings followed the order L-lysine, and 3 x glutamic acid. After final deprotection, a coupling was then performed in the same way with either then 4 eq. levulinic acid or **HT-ligand**. The resin was then washed with anhydrous DMF (3x10mL) and DCM (3x10mL) and transferred back to the PolyPrep® column. The DCM was then removed from the resin with positive $N_2$ pressure. Full cleavage and deprotection of the **CKEEE-HT** was accomplished with a solution of 95:2.5:2.5 TFA/$H_2O$/TIPS (20 mL, v/v/v). Full cleavage and deprotection of the **CKEEE-LA** was accomplished with a solution of 95:2.5:2.5 TFA:$H_2O$:TES (20 mL, v/v/v). The cleavage solutions were mixed via rotation for 120 min. The filtrate was then collected in a flask and the solvent was removed in vacuo. A small amount of oil with suspended white solids was obtained. Upon addition of diethyl ether (100 mL), white solids immediately crash out from the crude mixtures. The white solid was agitated in the ether and allowed to settle and the diethyl ether was decanted. The white solids were then washed in the same way an additional time and then dissolved in water (8 mL) and filtered through a 0.20 µm hydrophilic filters. The filtered solutions were frozen with liquid $N_2$ and lyophilized to yield flocculent white powders. Peptides were characterized ESI-MS and used without further purification (**Figure S3**). The cleavage of **CKEEE-LA** from resin lead to apparent reduction of the terminal ketone to an alcohol, likely by the TES. **CKEEE-HT**: ESI-MS: *m/z* calcd for

$C_{38}H_{65}ClN_7O_{16}S$: 942.39 $[M+H]^+$, found 942.6; **CKEEE-LA**: ESI-MS: *m/z* calcd for $C_{29}H_{49}N_6O_{14}S$: 737.30 $[M+H]^+$, found 737.4.

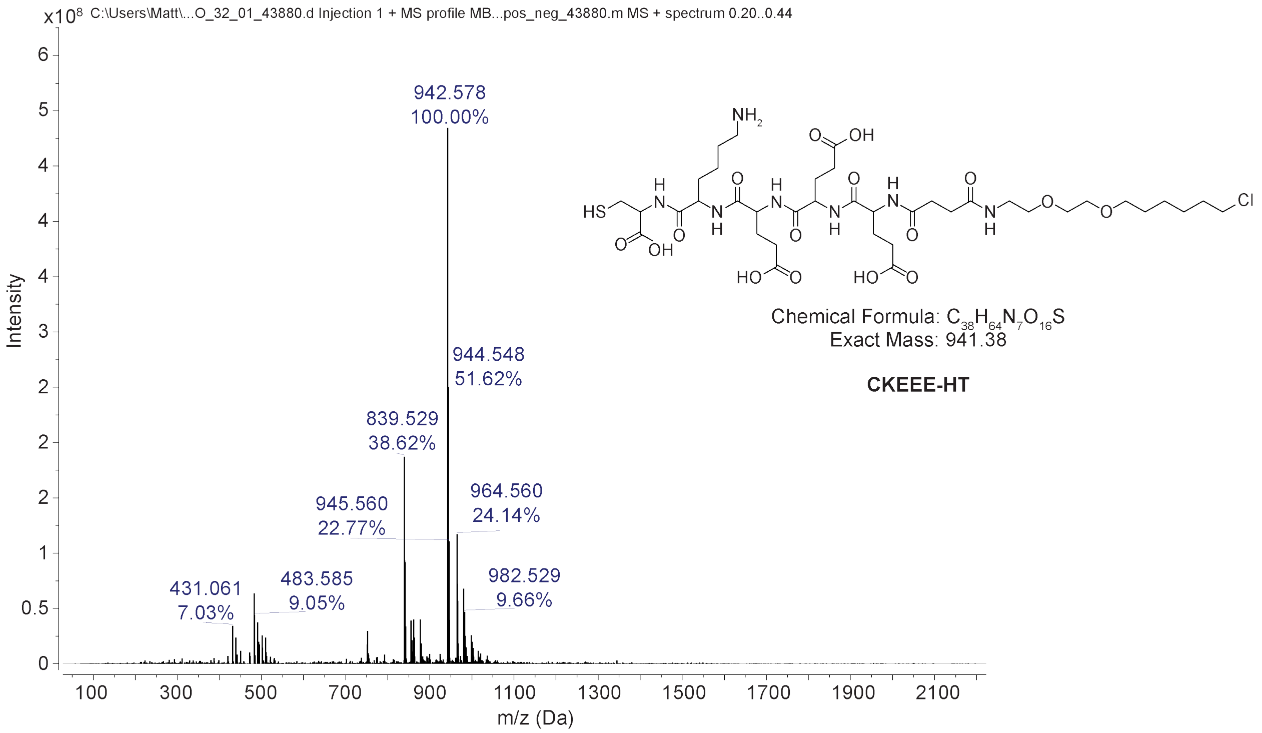


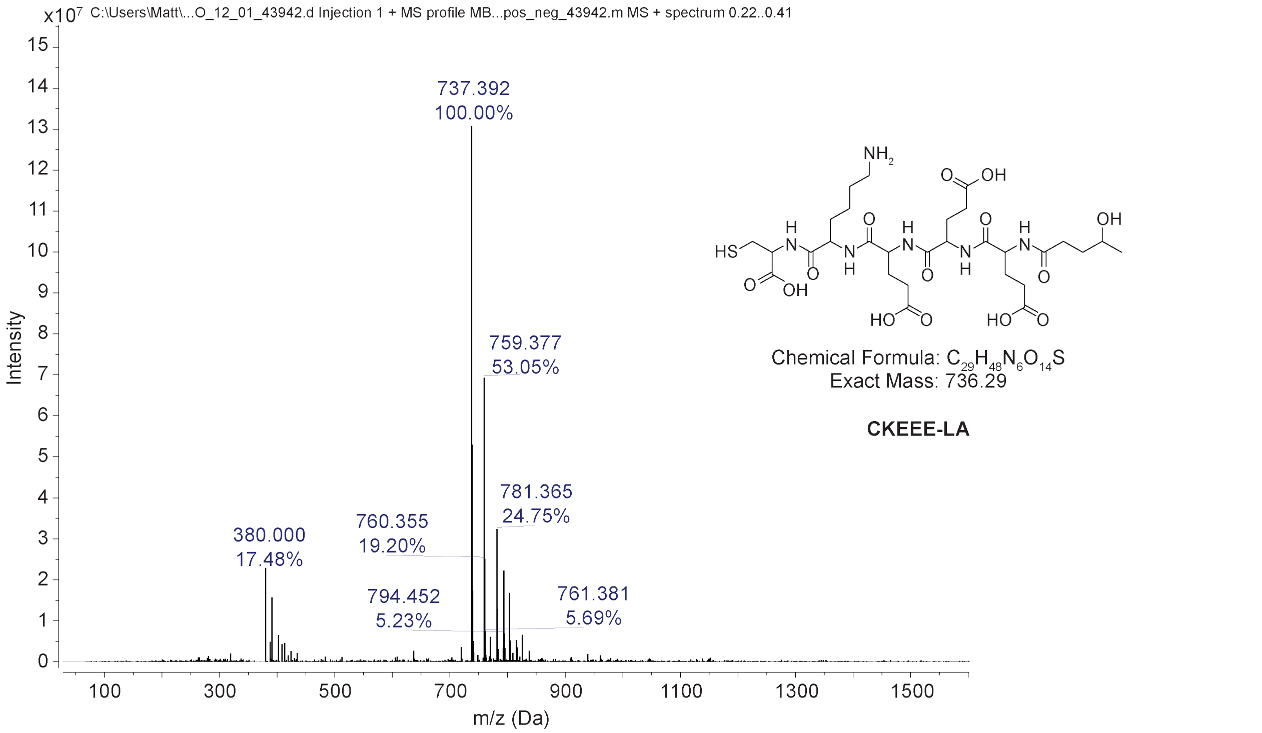


**Supplemental Figure 3:** Chemical structures, chemical formulae, calculated exact masses, and corresponding ESI-MS spectra of the synthetic **CKEEE-HT** (top) and **CKEEE-LA** (bottom) peptides.

**AuNP Functionalization Via Ligand Exchange**

Ligand exchanges were performed in water with undiluted colloid of citrate-stabilized colloidal AuNPs from synthesis. Two 5 mL aliquots of the gold colloid (calculated 23 mM concentration of AuNPs) were removed for ligand exchange. To each was added 6 µL of a 10% (v/v) TWEEN® 20 solution in water. Stock solutions of *ca.* 1 mM peptide concentration were made for both **CKEEE-HT** and **CKEEE-LA** by dissolving 1.0 mg of each into 1,060 µL and 1,360 µL of water, respectively. UV/vis was taken of each of these 1 mM peptide solutions (**Figure S4**). A mixture of 0.2 mM **CKEEE-HT** and 0.8 mM **CKEEE-LA** (1 mM total thiol) was made by combining 50 µL of CKEEE-HT stock solution with 200 µL of CKEEE-LA stock solution. To each aliquot of gold colloid was added either 150 µL of the **CKEEE-LA** or 20:80 **CKEEE-HT**/**CKEEE-LA** exchange solution (~1,300 molar equivalents of thiol per AuNP). There was no obvious change in the colloids upon exchange solution addition. Colloids were then rotated at low rpm for 16 h. To purify the free peptides away from conjugated AuNPs, the particles were pelleted via centrifuge (12,100 g, 30 min, 4°C) and 5 mL of supernatant was removed. To the remaining *ca.* 156 µL was added 5 mL fresh PBS and the pellet was re-suspended by vortexing. The particles were washed in the same way an additional time. The final colloid was then pelleted in the same way, 5 mL of supernatant was removed, and the pellet was re-suspended in only the remaining ~156 µL of PBS. Final colloid concentration is estimated at 0.7 µM functionalized AuNPs.

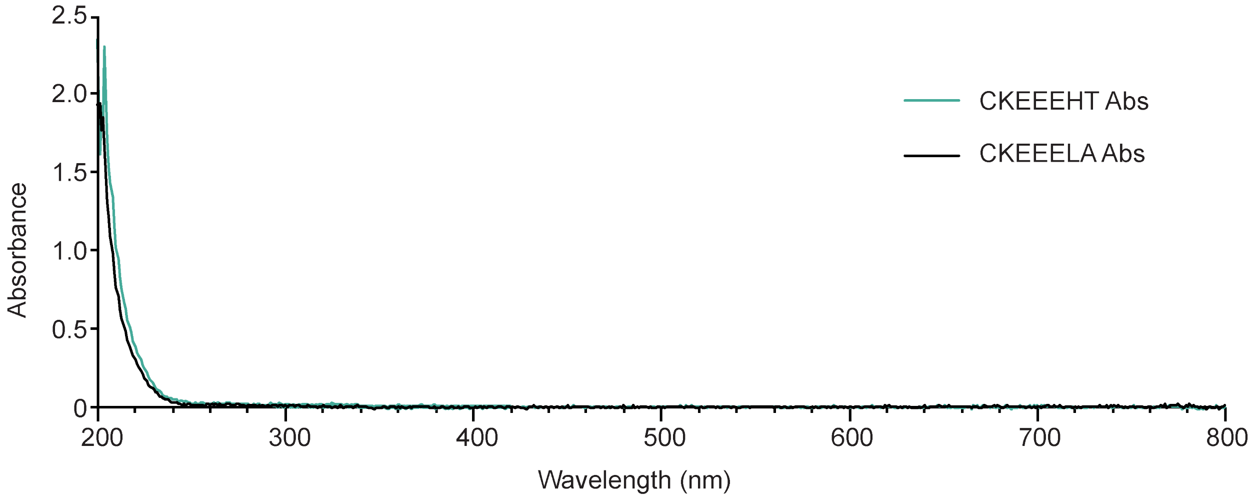


**Supplementary Figure 4**: UV/visible spectra of *ca.* 1 mM solutions of **CKEEE-HT** and **CKEEE-LA** used for ligand exchange reactions AuNPs collected with a 1 mm path length.

#### ADDITIONAL SUPPLEMENTARY FIGURES

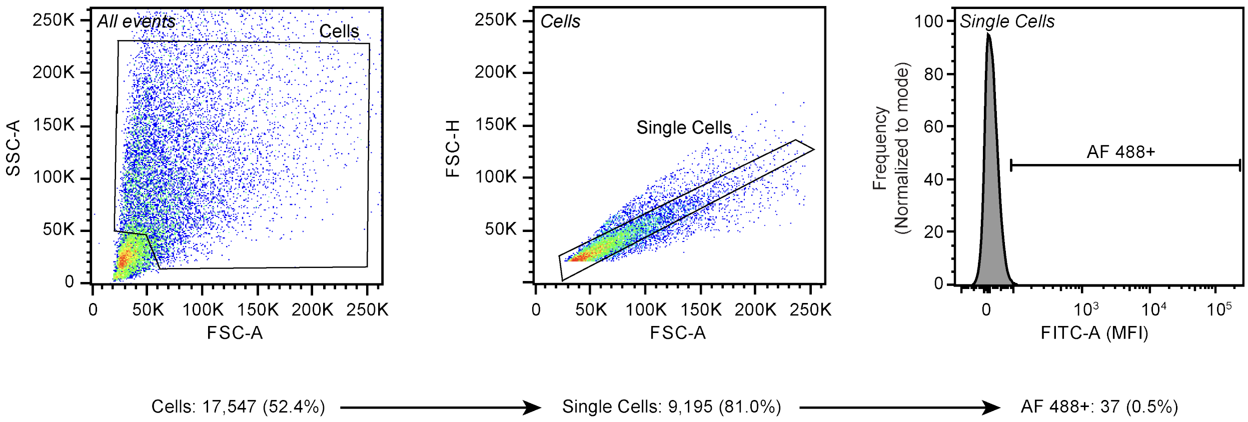


**Supplemental Figure 5:** Conventional flow cytometry gating scheme for cells. The plots show a sample of unmodified cells. These cells do not express HaloTag and were not labeled with the AF 488 HaloTag ligand. In the gating procedure, cells were identified based on the FSC-A vs SSC-A profile. From this population, single cells were identified based on the FSC-A vs FSC-H profile. The AF 488+ cell population was defined as all single cells with a greater FITC signal than the sample of unmodified cells. This gate was drawn such that it did not encompass more than 0.5% of this non-fluorescent population of cells.

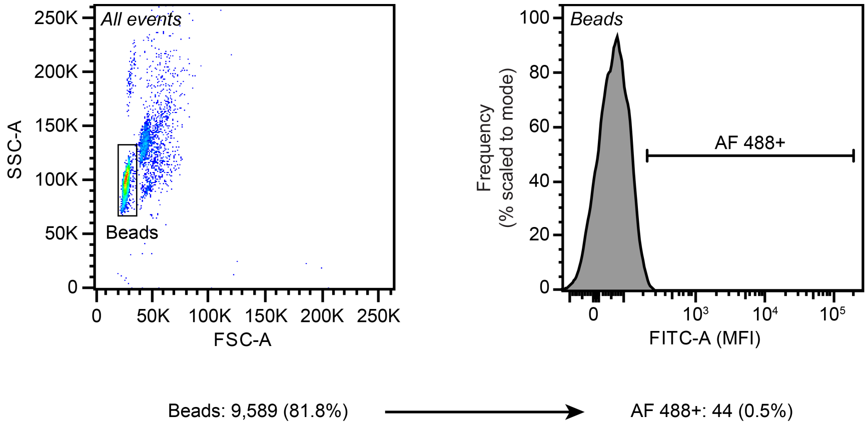


**Supplemental Figure 6:** Conventional flow cytometry gating scheme for EVs adsorbed on beads. The plots show a sample of unmodified EVs adsorbed onto latex beads. These EVs are not loaded with HaloTag and were not labeled with the AF 488 HaloTag ligand. In the gating procedure, beads were identified based on the FSC-A vs SSC-A profile. The AF 488+ bead population was defined as all beads with a greater FITC signal than beads adsorbed with EVs harvested from unmodified cells. This gate was drawn such that it did not encompass more than 0.5% of this non-fluorescent population of beads.

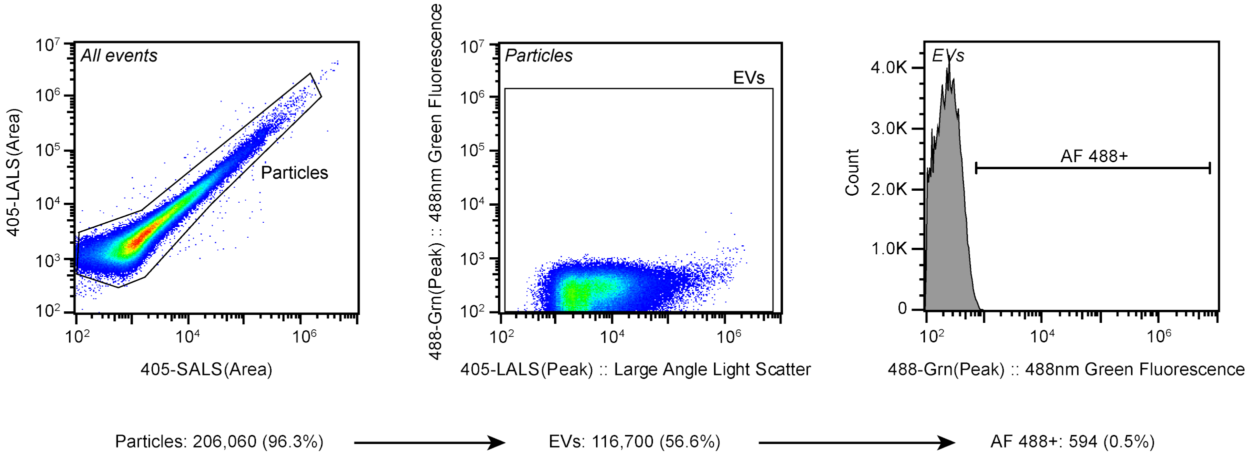


**Supplemental Figure 7:** Apogee single vesicle flow cytometry gating scheme for EVs. The plots show a sample of unmodified MVs run on an Apogee single vesicle flow cytometer. These EVs were not loaded with HaloTag and were not labeled with the AF 488 HaloTag ligand. In the gating procedure, particles were identified based on the 405-SALS (Area) vs 405-LALS (Area) profile. From this population, EVs were identified based on the 405-LALS (Peak) vs 488-Grn(Peak) profile. The AF 488+ EV population was defined as all EVs with a greater 488 nm green fluorescence signal than the sample of EVs harvested from unmodified cells. This gate was drawn such that it did not encompass more than 0.5% of this non-fluorescent population of EVs.

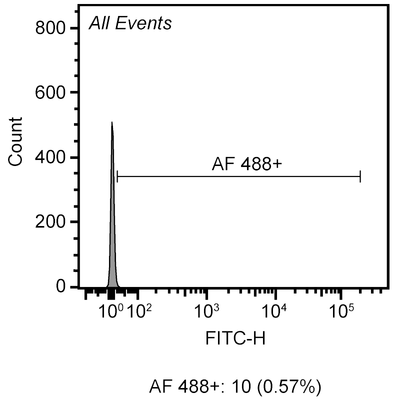


**Supplemental Figure 8:** NanoFCM single vesicle flow cytometry gating scheme for EVs. The plot show a sample of exosomes run on an NanoFCM single vesicle flow cytometer. The AF 488+ EV population was defined as all EVs with a 488 nm green fluorescence signal greater than the background sample (unmodified EVs treated with AF 488 ligand). This gate was drawn such that it did not encompass more than 0.5% of this non-fluorescent sample, to account for ligand remaining after EV conjugation and washing. The same gating strategy was applied for AF 660 conjugated EVs, using unmodified EVs treated with AF 660 ligand as the background sample.

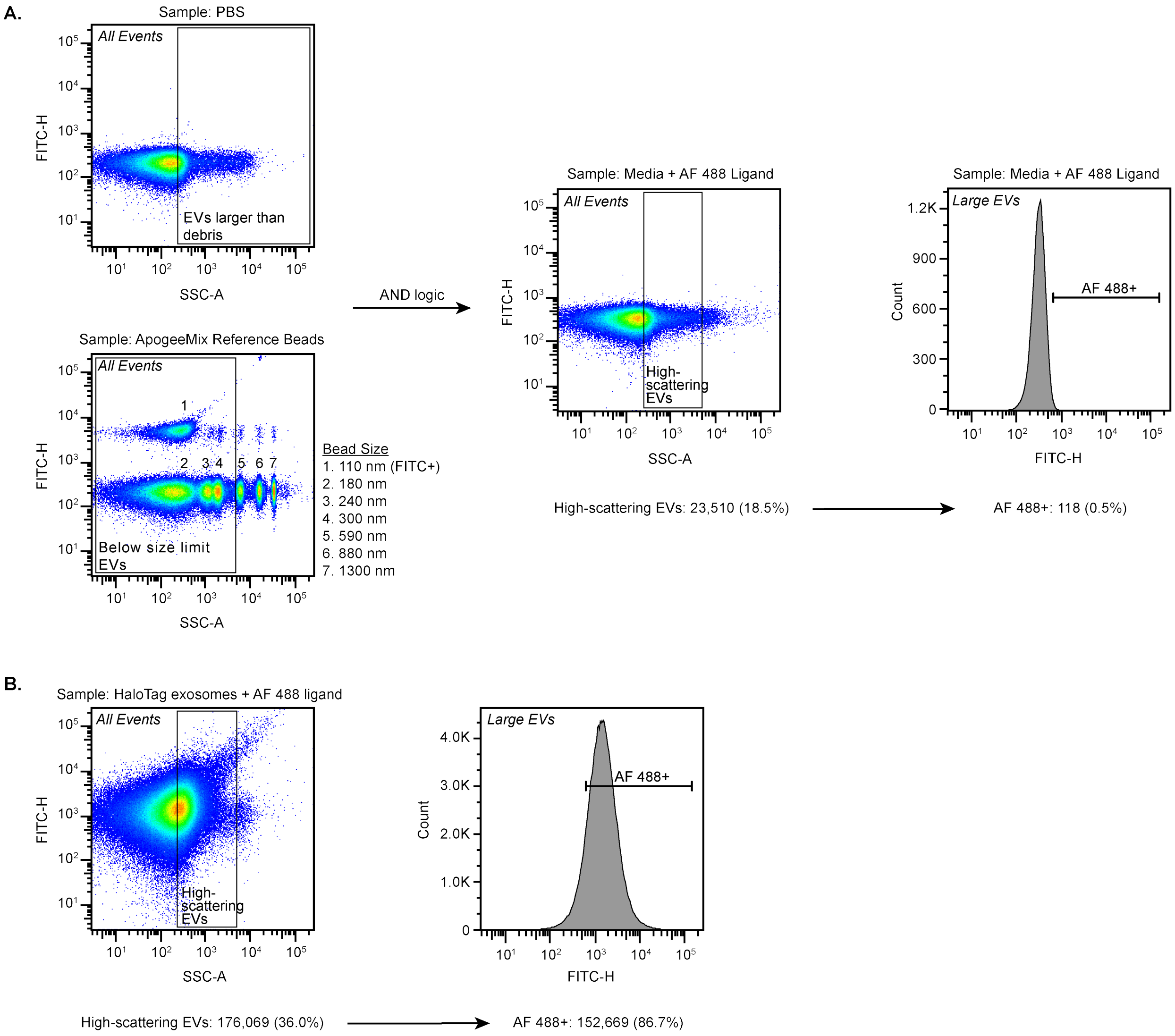


**Supplemental Figure 9:** Conventional BD LSR Fortessa flow cytometer gating scheme for single EVs. A) In the gating procedure for AF 488 conjugated EVs, a sample of 0.1 um filtered PBS on a FITC-H vs SSC-A plot was used to exclude small events which we considered non-EV background debris and bubbles present in all samples. EVs in the size range of interest (<590 nm) were identified using ApogeeMix Size reference beads (Apogee, #1527) on a FITC-H vs SSC-A plot. AND logic was used to combine these gates, resulting in a population termed "Large EVs" that is enriched in large EVs in the size range of interest. The AF 488+ EV population was defined as all EVs with a greater 488 nm green fluorescence signal than the mock sample (media treated with AF 488 ligand). This gate was drawn such that it did not encompass more than 0.5% of this non-fluorescent sample, to account for ligand remaining after EV conjugation and washing. The same gating strategy was applied for AF 660 conjugated EVs, using media treated with AF 660 ligand as the mock sample. B) Example of gating on a sample of HaloTag exosomes labeled with AF 488 ligand, resulting in 86.7% of exosomes being identified as AF 488+.

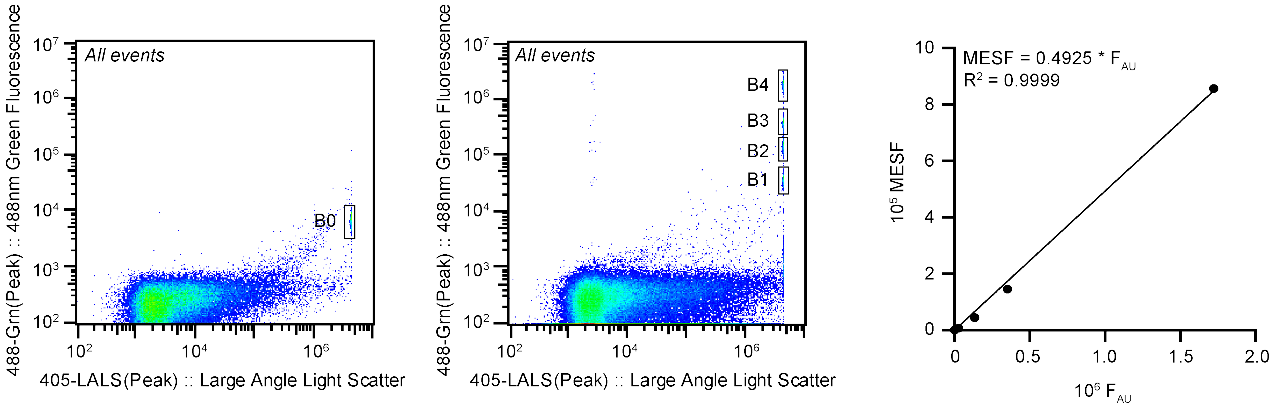


**Supplemental Figure 10:** Apogee Micro Plus profile of fluorescence quantification beads. Bangs Lab Quantum™ Alexa Fluor 488 MESF beads have 4 fluorescent bead populations (B1-4) and 1 blank population (B0). Beads were identified based on the 405-LALS (Peak) vs 488-Grn(Peak) profile. The mean intensity of each bead population in the 488-Grn(Peak) channel in arbitrary units ($F_{AU}$) was recorded, the mean intensity of B0 was subtracted, and the resulting background subtracted $F_{AU}$ was plotted against the manufacturer-supplied number of fluorophores on the beads for each population (MESF). To generate the calibration curve, a linear regression was performed with the constraint that the y-intercept equals zero. A new calibration curve was generated for each experiment, and EV MFI was converted to MESF by using the multiplier on $F_{AU}$ obtained from the regression.

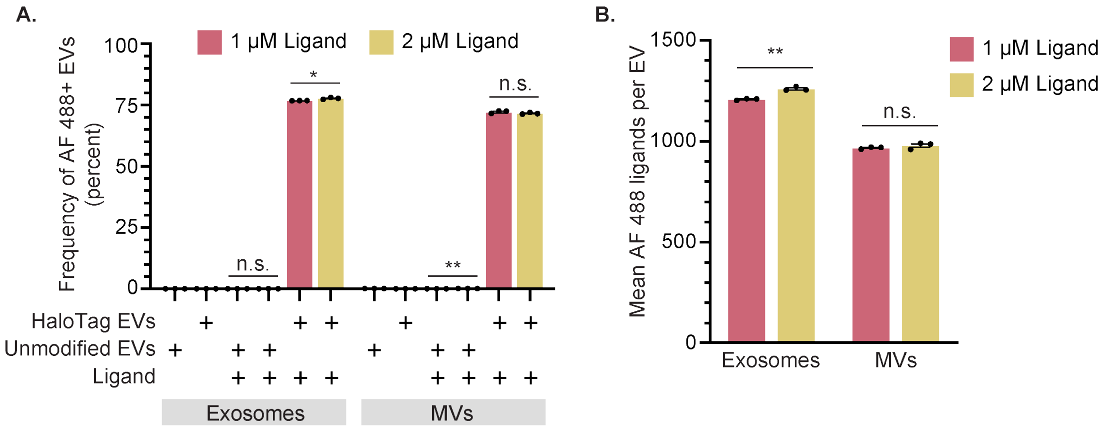


**Supplemental Figure 11:** HaloTag EV conjugation reaction proceeds to completion at manufacturer's recommendation of 1 µM AF 488 ligand concentration. A) The frequency of AF 488+ EV labeling changes minimally when the ligand reaction concentration is increased to 2 µM compared to the 1 µM; all analyses employ Apogee SV-FC. B) Degree of AF 488 ligand conjugated per EV varies moderately for exosomes and negligibly for MVs when excess ligand is present. EV experiments were performed in technical triplicate. Error bars indicate standard error of the mean. Pairwise comparisons were made using unpaired Student's t-test. Multicomparison statistical analysis was performed using a one-way ANOVA test, followed by Tukey's multiple comparison test to evaluate specific comparisons (*$p < 0.05$, **$p < 0.01$, ***$p < 0.001$, ****$p < 0.0001$).

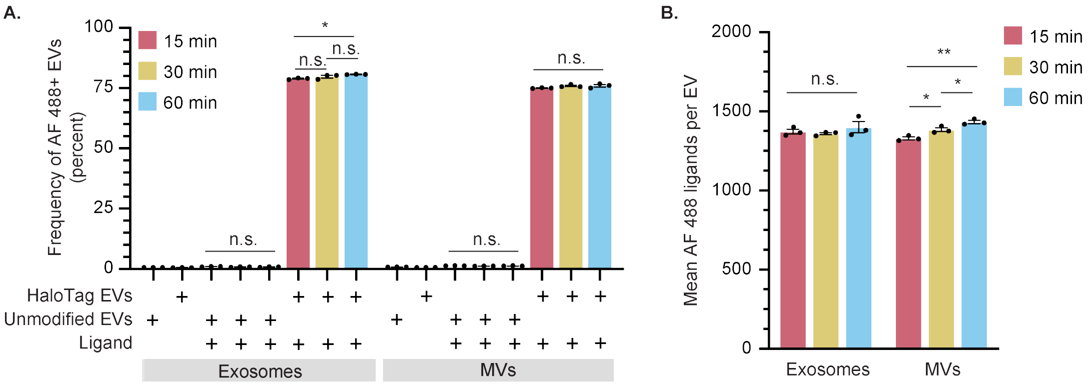


**Supplemental Figure 12:** The HaloTag EV conjugation reaction proceeds to completion in 15 minutes. A) Increase in the percent of AF 488+ EVs is minimal between 15-60 minutes; all analyses employ Apogee SV-FC. B) Increase in the amount of AF 488 ligand conjugated per EV is negligible for exosomes and moderate for MVs. EV experiments were performed in technical triplicate. Error bars indicate standard error of the mean. Pairwise comparisons were made using unpaired Student's t-test. Multicomparison statistical analysis was performed using a one-way ANOVA test, followed by Tukey's multiple comparison test to evaluate specific comparisons (*$p < 0.05$, **$p < 0.01$, ***$p < 0.001$, ****$p < 0.0001$).

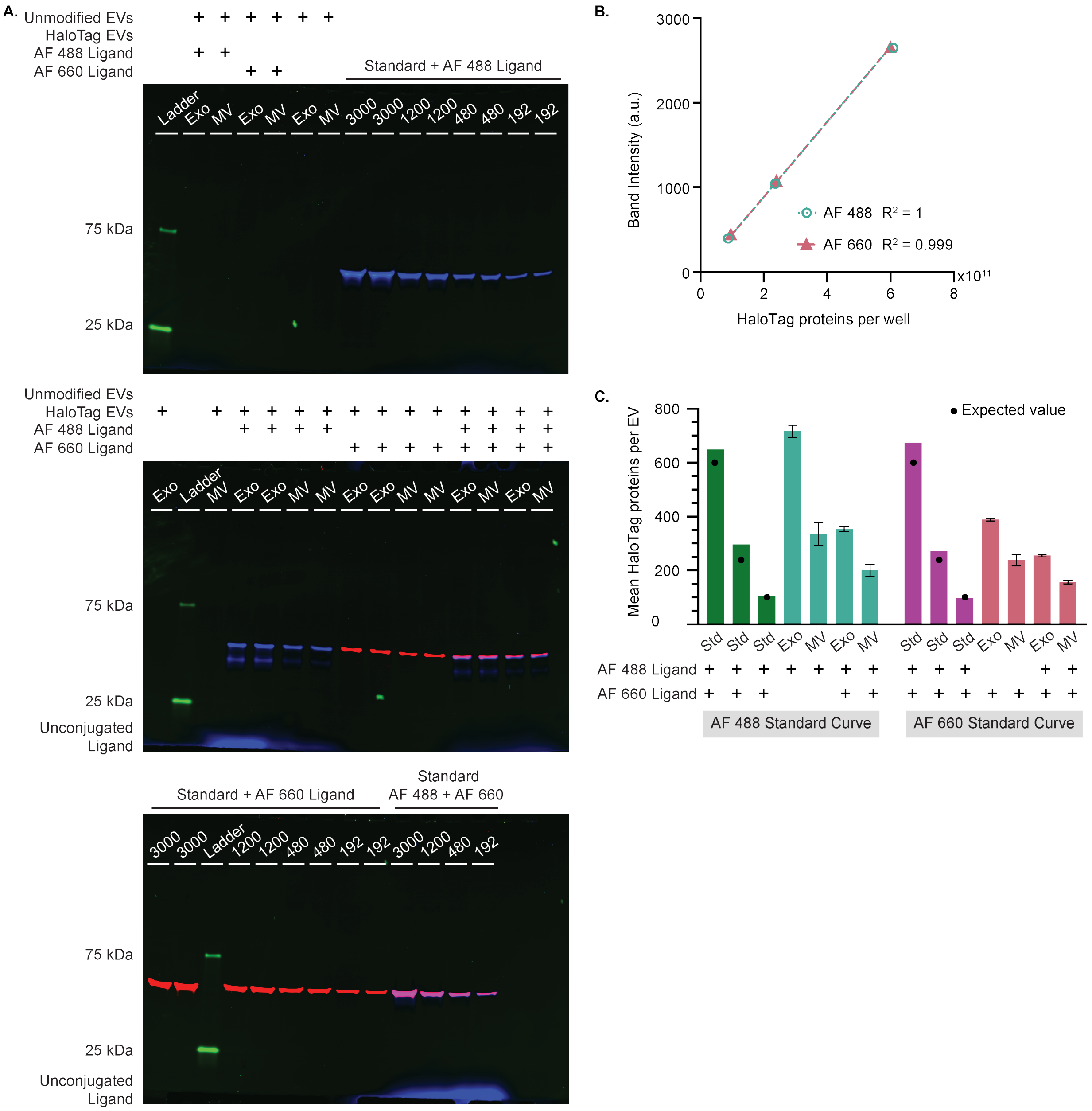


**Supplementary Figure 13:** Protein gels and standard curves show functional HaloTag construct loading into EVs. A) Composite fluorescent image with signal from the 488 nm laser colored blue, 520 nm laser colored green, and 658 nm laser colored red. HaloTag EVs and HaloTag standard conjugated with AF 488 and AF 660 are blue and red, respectively. Co-labeled EVs are purple. EVs with no HaloTag protein or no conjugated ligand have no signal in any channel. Protein ladder shows the expected 25 and 75 kDa bands in green. Both EV populations contain the expected band size of 44.8 kDa. Values at the top of the standard lanes indicate the number of recombinant proteins added to each well divided by 4.5 x $10^8$ (the number of vesicles added to the other wells). Blue band present at the bottom of the gel images is unconjugated AF 488 dye. B) Mean fluorescent band intensity of lane was calculated and plotted against the amount of HaloTag standard protein loaded. To generate the calibration curve, a linear regression was performed with the constraint that the y-intercept equals zero. C) The average amount of HaloTag proteins per EV was calculated using the mean fluorescent band intensity of the lane and the corresponding ligand’s standard curve. Co-labeled samples contained both ligands at the recommended reaction concentrations such that there should be equal labeling of both ligands. Co-labeled protein standards are labeled with the expected equivalent mean HaloTag proteins per EV, showing good agreement with measured results. The expected 50% reduction when co-labeled is observed for both exosomes and MVs when quantifying with the AF 488 standard curve. When using the AF 660 calibration curve, there are fewer HaloTag proteins per EVs for both EV populations compared to the AF 488 calibration curve, and there is a smaller reduction in HaloTag proteins measured per EV for co-labeled EVs.

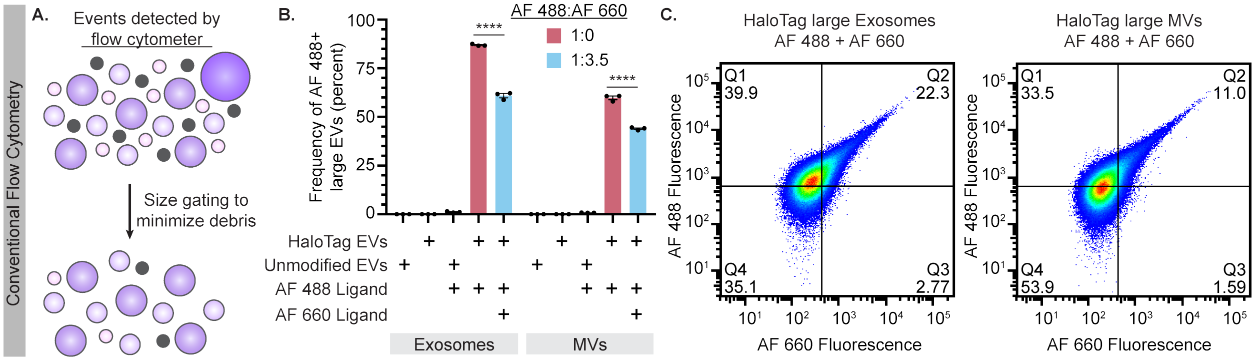


**Supplemental Figure 14:** A-C) Analysis of EVs by conventional flow cytometry. A) This cartoon illustrates events detected by a conventional flow cytometer, comprised of EVs of various sizes (purple) and bubbles and debris (gray) which overlaps in size with small EVs (~100 nm). Conservative size gating was applied to minimize non-EV events including events larger than the size of EVs of interest (>600 nm), resulting in a population enriched in "large EVs". B) Frequency of AF 488+ large exosomes measured on a conventional flow cytometer agrees well with data collected via Apogee SV-FC, whereas frequency of AF 488+ large MVs is lower than observed by Apogee. Frequency of AF 488+ large EVs decreases significantly with increasing AF 660 ligand concentration. C) HaloTag EVs can be co-labeled to be AF 488+ and AF 660+ (double-positive). EV experiments were performed in technical triplicate. Error bars indicate standard error of the mean. Pairwise comparisons were made using unpaired Student's t-test. Multicomparison statistical analysis was performed using a one-way ANOVA test, followed by Tukey's multiple comparison test to evaluate specific comparisons (*$p < 0.05$, **$p < 0.01$, ***$p < 0.001$, ****$p < 0.0001$).

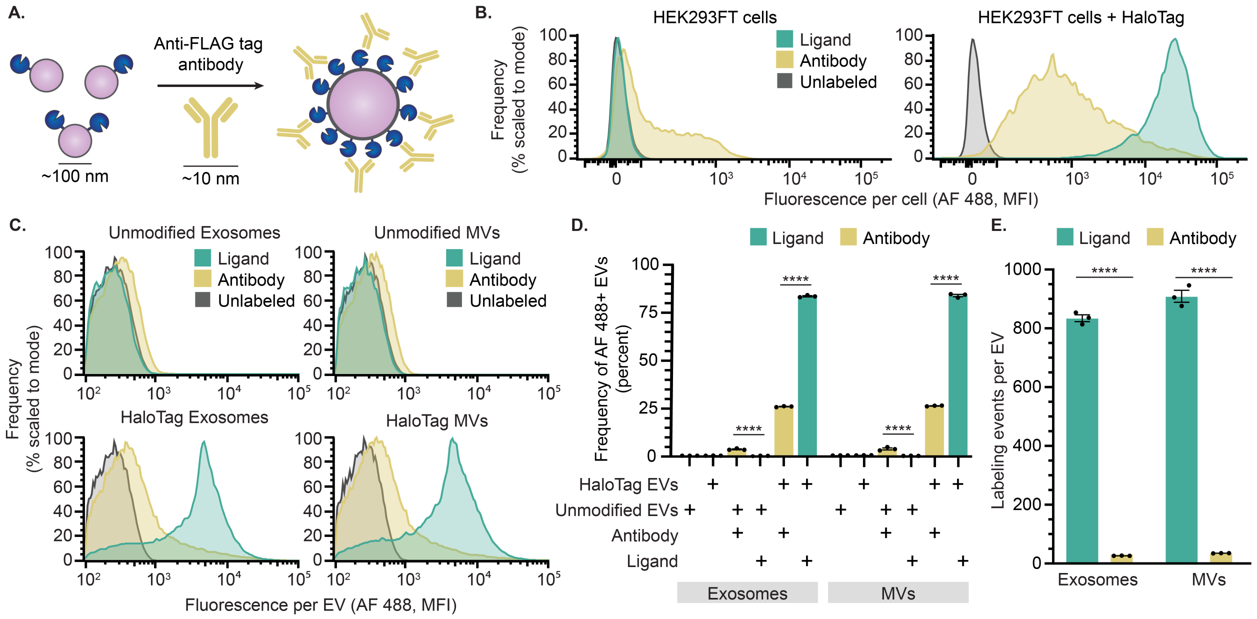


**Supplemental Figure 15:** Anti-FLAG tag antibody labeling of HaloTag EVs underestimates protein display. A) This cartoon illustrates anti-FLAG tag antibody-mediated labeling of HaloTag protein on EVs. HaloTag protein (~4 nm) and EVs (~100 nm) are not to scale. B) Antibody or HaloTag ligand-mediated labeling of EV producer cells. C) Antibody labeling of EVs compared to HaloTag conjugation; all analyses employ Apogee SV-FC. D) Antibody labeling underestimates the frequency of HaloTag-expressing EVs relative to HaloTag-conjugation. E) Antibody labeling of HaloTag EVs underestimates the number of HaloTag proteins per EV. Cell experiments were performed in biological triplicate. EV experiments were performed in technical triplicate. Error bars indicate standard error of the mean. Pairwise comparisons were made using unpaired Student's t-test. Multicomparison statistical analysis was performed using a one-way ANOVA test, followed by Tukey's multiple comparison test to evaluate specific comparisons (*p < 0.05, **p < 0.01, ***p < 0.001, ****p < 0.0001).

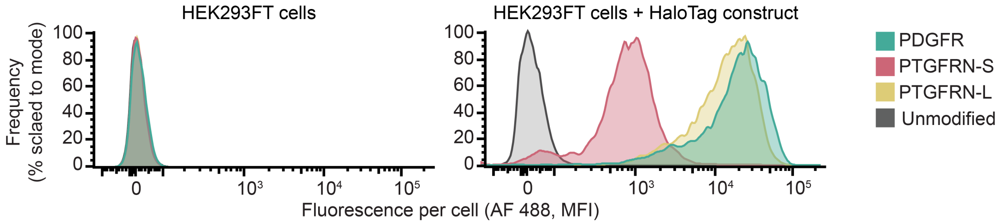


**Supplemental Figure 16:** PTGFRN-S TMD display construct yields lower levels of HaloTag display on cells compared to PDGFR TMD and PTGFRN-L TMD display constructs. Cells were conjugated with AF 488 ligand, washed, and analyzed by flow cytometry.

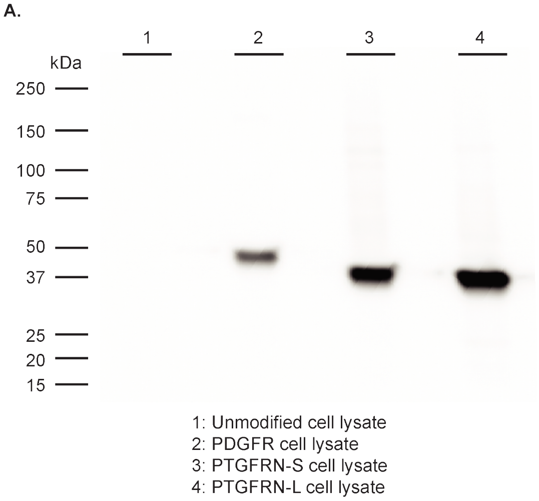


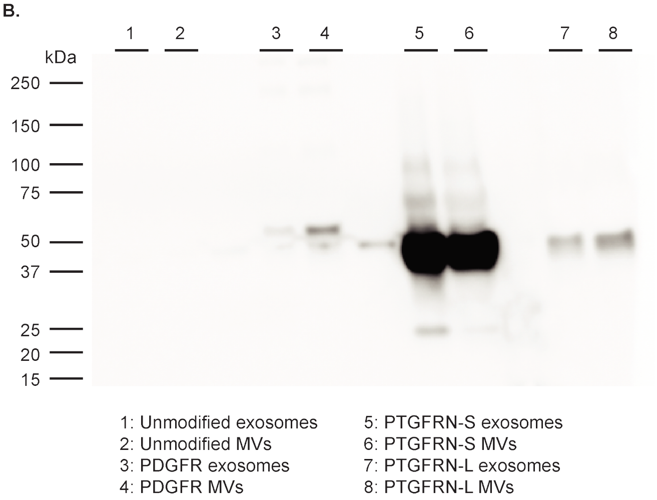


**Supplemental Figure 17:** HaloTag fusion proteins are detected in cell lysates and in both EV populations when detected with anti-Flag tag antibody (Flag tag is fused to N-terminus of engineered HaloTag). A) Cell lysates show the expected band sizes: PDGFR - 44.8 kDa; PTGFRN-S – 41.5 kDa; PTGFRN-L – 42.4 kDA. B) Both EV populations contain the expected band size for all three fusion proteins. PTGFRN-S EVs are loaded with significantly more protein than PDGFR and PTGFRN-L EVs.
